# Stress Granule Formation Drives ZBP1-dependent Necroptosis in Non-obstructive Azoospermia and Aging testes

**DOI:** 10.1101/2025.04.11.648478

**Authors:** Hongen Lei, Dianrong Li, Jie Chen, Baowen Du, Kaiju Jiang, Tao Xu, Hu Han, Weiliang Fan, Long Tian, Xiaodong Wang

**Author notes:** Corresponding author. Email: Dianrong. These authors contributed equally (Hongen Lei, Dianrong Li).

## Abstract

**Male infertility** and **age-related reproductive decline** remain major unmet medical challenges, with limited understanding of the underlying mechanisms. Here, we identify a **stress granule–driven necroptosis pathway** involving **ZBP1 and RIPK3** as a central driver of **non-obstructive azoospermia (NOA)**, a severe form of male infertility marked by loss of spermatogenesis. We show that **heat stress** or environmental insults activate **eIF2α kinases**, inducing stress granules that recruit **ZBP1 and RIPK3** to form a signaling complex. This leads to **RIPK3 activation**, **MLKL phosphorylation**, and necroptotic death of **spermatogonia and Sertoli cells**. Genetic ablation of ***Zbp1*** or ***Ripk3*** protects mice from **heat-induced testicular atrophy**, highlighting their essential role in testicular cell death. **Notably**, this same necroptosis pathway is also activated in **aged human testes**, suggesting a shared mechanism driving both **male infertility** and **age-related testicular degeneration.**

## Introduction

**Male infertility** affects up to 15% of couples worldwide, with azoospermia— complete absence of sperm in semen—among its most severe forms^1,2^. Non-obstructive azoospermia (NOA), characterized by testicular failure, remains poorly understood at the molecular level despite its clinical prevalence^3,4^. Testicular aging similarly leads to diminished spermatogenesis^5–7^, yet the cellular processes linking infertility and aging remain elusive.

**Necroptosis**, a regulated form of inflammatory cell death, has emerged as a common feature of neurodegeneration, viral infection, and tissue injury, but its relevance to infertility is unknown^8–10^. We previously observed activation of necroptosis in spermatogonia and Sertoli cells of aged mice, along with phospho-MLKL enrichment in aged human testes^11,12^. Here, we demonstrate that **NOA is driven by stress granule–induced necroptosis** mediated through the **ZBP1–RIPK3–MLKL pathway**. We show that environmental stress triggers eIF2α kinase activation and stress granule assembly, which recruits ZBP1 and RIPK3 to initiate necroptosis. Testicular biopsy samples from NOA patients exhibited robust phospho-MLKL and stress granule markers in all cases examined. Genetic deletion of *Zbp1* or *Ripk3* prevented heat-induced testicular atrophy in mice, confirming the essential role of this pathway. **Notably, we also detect activation of this pathway in aged human testes**, revealing a shared mechanism linking infertility and reproductive aging. These findings identify stress granule–mediated necroptosis as a key contributor to male reproductive decline and suggest new molecular targets for intervention.

## Results

### Patient Cohort and Testicular Histopathology

We recruited 50 unrelated patients diagnosed with **non-obstructive azoospermia (NOA)** for histological and molecular analyses. Diagnosis was based on **AUA/ASRM clinical guidelines**^13^, and the final cohort included 40 cases of **idiopathic NOA** and 10 with a history of **cryptorchidism** (Extended Data Fig. 1). Control testicular samples were obtained from 17 patients undergoing **orchiectomy for testicular torsion**, characterized by ischemia-related loss of blood flow. Histological evaluation revealed that **NOA testes lacked sperm** and showed a **Johnsen Score ≤ 7**, indicative of **complete spermatogenic failure**, while torsion samples retained normal spermatogenesis (Fig. 1a–c, Extended Data Fig. 2, Supplementary Table 1). The mean age of NOA patients at diagnosis was 32 years.

**Fig. 1.**
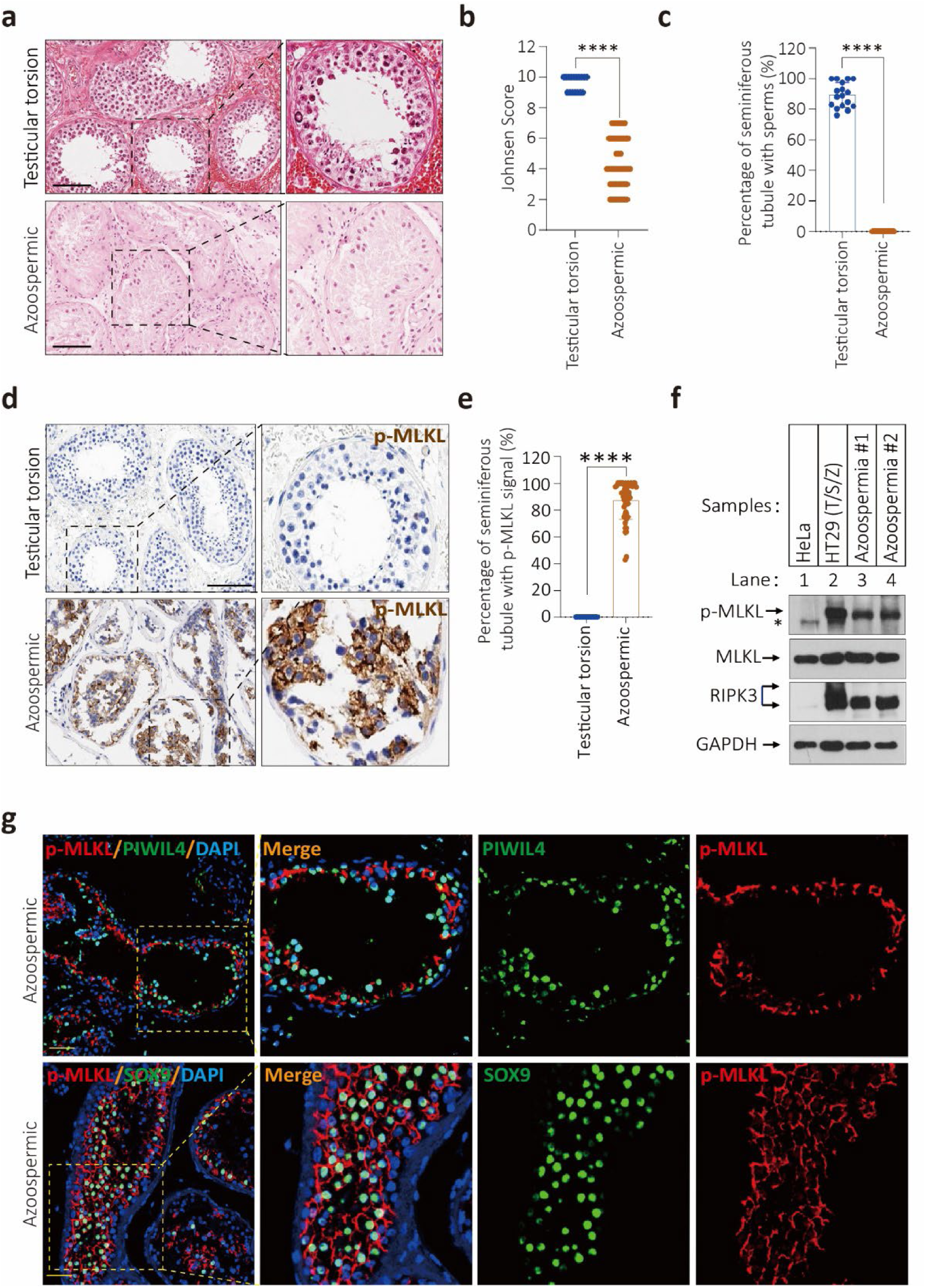
Phosphor-MLKL(p-MLKL) were detected in the seminiferous tubules of NOA patients’ testes. **a**-**c**, Hematoxylin and Eosin (H&E) staining of testis sections from human testicular torsion(n=17) and azoospermia patients(n=50) in (a). Johnsen Score evaluation of testicular torsion and azoospermia patients based on (a) and show in (b). The number of seminiferous tubules with sperm were counted based on (a) and quantification in (c), seminiferous tubules with sperm were counted in five fields per testis. Scale bar, 100 μm. **d**, **e**, Immunohistochemistry (IHC) analysis of human testicular torsion and NOA testis sections with p-MLKL antibody in (d). The number of seminiferous tubules with positive p-MLKL signal were counted based on IHC staining and quantification in (e). Scale bar, 100 μm. **f**, Western blotting analysis of RIPK3, p-MLKL and MLKL in the testis from two NOA patients, GAPDH was used as loading control. HeLa and HT29 cells treated with the combination of T/S/Z as negative and positive control for western blotting analysis respectively. The asterisk(*) indicates non-specific bands. **g**, Immunofluorescence analysis of NOA testis sections with antibodies against p-MLKL (red) PIWIL4 (spermatogonium specific protein, green) and SOX9 (Sertoli cells specific protein, green). Scale bar, 50 μm. Quantified data in (b, c and e) represent the mean ± s.e.m. \*\*\*\**P*<0.0001. *P* values were determined by two-sided unpaired Student’s *t* tests.

### Phospho-MLKL Is Robustly Induced in NOA Testes

Given prior evidence linking **phospho-MLKL – driven necroptosis** to aging-related testicular decline in mice and humans^11,12,14^, we investigated whether this pathway was similarly activated in NOA. **Immunohistochemistry (IHC)** using a phospho-MLKL–specific antibody revealed **robust signal in the seminiferous tubules** of all NOA patients (100%, n = 50)**, but** no staining in torsion controls (0%, n = 17) (Fig. 1d–e). To confirm antibody specificity, we performed two independent controls: (1) peptide competition with phosphorylated MLKL peptides, and (2) **lambda protein phosphatase (LPP)** pretreatment. Both eliminated the IHC signal, validating the specificity of phospho-MLKL detection (Extended Data Fig. 3a–b).

### RIPK3, Not RIPK1, Drives MLKL Activation in NOA

As **RIPK3** is the only known kinase that phosphorylates MLKL, its activity is inferred by phospho-MLKL presence^8–10,15^. We next asked whether RIPK3 was activated via canonical RIPK1-dependent signaling. **Phospho-RIPK1** was undetectable in all NOA samples (0%, n = 50), indicating that **RIPK3 activation in NOA occurs independently of RIPK1** (Extended Data Fig. 4a). **Western blotting** of testis lysates from two NOA patients confirmed the presence of phospho-MLKL. Positive and negative controls included HeLa and HT29 cells treated with TNF-α, Smac mimetic, and the pan-caspase inhibitor Z-VAD-fmk (T/S/Z) (Fig. 1f). **Cleaved caspase-3**, a marker of apoptosis, was undetectable in all NOA samples (0%, n = 50), further supporting **necroptosis** as the dominant cell death pathway in NOA (Extended Data Fig. 4b).

### Necroptosis Occurs in Spermatogonia and Sertoli Cells

To identify the cell types undergoing necroptosis, we performed IHC co-staining using **cell-type-specific markers**. Phospho-MLKL co-localized with **PIWIL4** (spermatogonia) and **SOX9** (Sertoli cells), and to a lesser extent with **DDX4** (spermatocytes/spermatids) (Fig. 1g, Extended Data Fig. 5). In contrast, **Leydig cells**, which reside outside the seminiferous tubules and lack RIPK3 expression, did not exhibit phospho-MLKL signal.

### Heat Stress Induces ZBP1–RIPK3–Dependent Necroptosis

The testes are positioned outside the body to maintain a temperature **2–4°C below core body temperature**, a requirement critical for **spermatogenesis**^16,17^. In our NOA patient cohort, we noted that the **10 individuals with cryptorchidism**—a condition in which one or both testes fail to descend out of the body—exhibited **stronger phospho-MLKL staining** than those with idiopathic NOA, suggesting that **elevated testicular temperature may promote necroptosis**. Given prior reports implicating **ZBP1 in heat stroke–induced necroptosis**^18^, we examined whether **heat shock (HS)** alone could induce necroptotic death in mouse testicular cell lines. Exposure of **GC-2spd(ts)** (spermatocytes), **15P-1** (Sertoli cells), and **MA-10** (Leydig cells) to **43°C** did not elicit substantial cell death (Fig. 2a–b), nor did it in **L929 mouse fibroblast cells**, even 24 hours post-heat shock (Extended Data Fig. 6a–b). Because **ZBP1 expression is known to be IFN-β–inducible**^19,20^, we pretreated those cells with **interferon-β** before heat shock. Strikingly, IFN-β–primed GC-2spd, 15P-1, and L929 cells exhibited **significant cell death** upon heat stress, whereas vehicle (DMSO)-treated controls did not (Fig. 2b, Extended Data Fig. 6b–c). These results indicate that **IFN-β–induced gene expression is required for heat-induced necroptosis**. Consistent with this, **phospho-MLKL (Ser345)**—the murine equivalent of human Ser358—was readily detected in GC-2spd and 15P-1 cells after **IFN-β/HS treatment** (Fig. 2c). In contrast, **MA-10 cells**, which lack **RIPK3 expression**, failed to show either **cell death or phospho-MLKL** signal following IFN-β/HS exposure (Fig. 2b–c), confirming **RIPK3 dependence**.

**Fig. 2.**
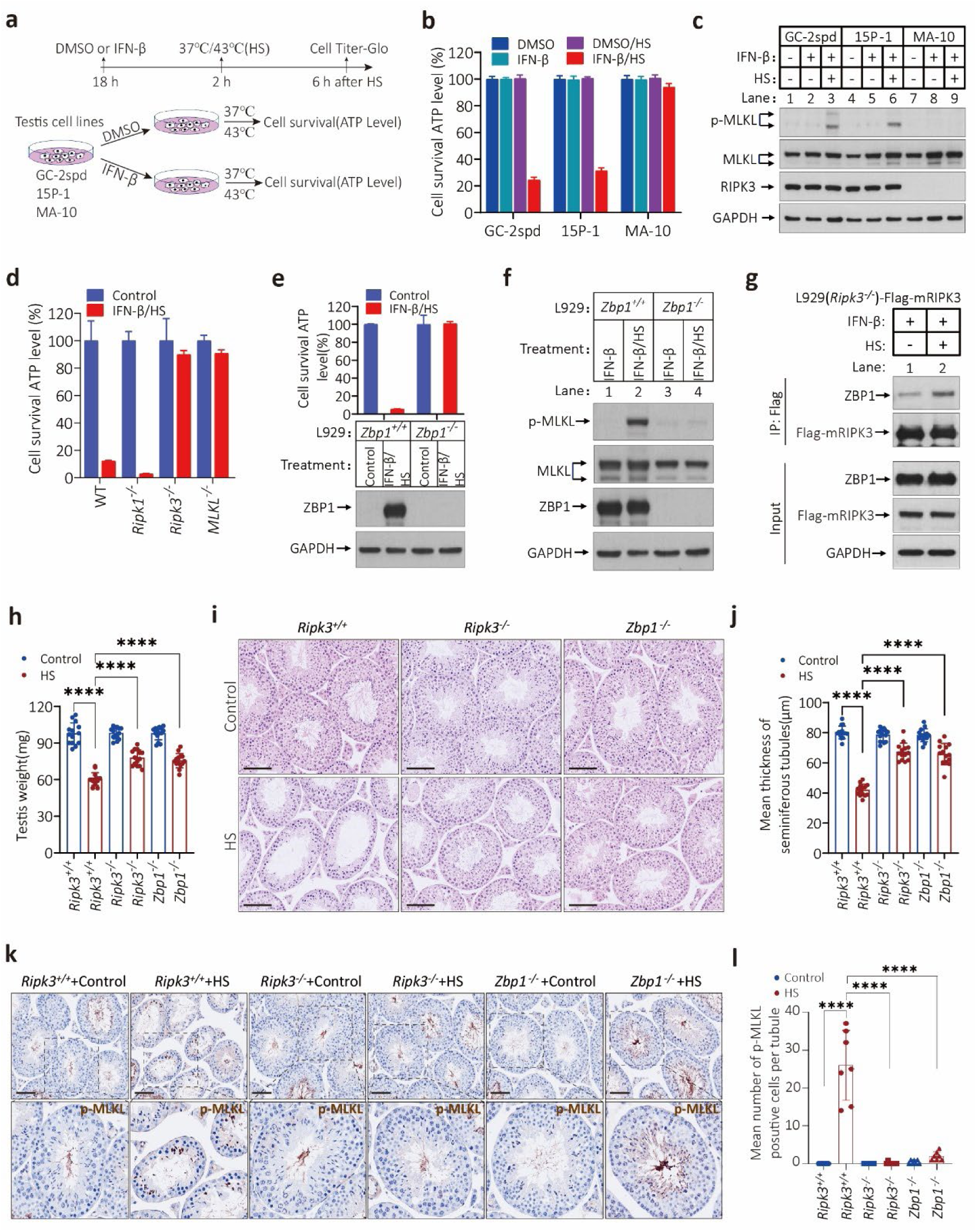
Heat shock-induced necroptosis dependents on ZBP1 and RIPK3. **a**-**c**, Schematic illustrating experiment design in (a). Cultured GC-2spd, 15P-1 and MA-10 cells were treated with DMSO or IFN-β for 18 hours, then the cells were transferred to 1.5 ml Eppendorf Tubes (EP) and put into 37℃ or 43℃ water for 2 hours. 6 hours after heat shock (HS) cell viability as measured by Cell Titer-Glo in (b). The levels of p-MLKL, MLKL and RIPK3 were analyzed by immunoblotting in (c), GAPDH was used as loading control. **d**, Cultured L929 cells with wild type (WT), *Ripk1*, *Ripk3* or *Mlkl* gene knocked out were treated with IFN-β and HS for 18 hours and 1.5 hours as described in Extended Data Fig. 6a, 2 hours after heat shock cell viability as measured by Cell Titer-Glo. **e**, **f**, Cultured L929 cells with wildtype or *Zbp1* gene knocked out were treated with IFN-β and HS for 18 hours and 1.5 hours as described in Extended Data Fig. 6a, 2 hours after heat shock cell viability as measured by Cell Titer-Glo in (e). The levels of p-MLKL, MLKL, and ZBP1 were analyzed by immunoblotting in (f), GAPDH was used as loading control. **g**, Cultured L929(*Ripk3^-/-^*)-HA-3×Flag-mRIPK3 cells were treated with IFNβ/HS as indicated. The cell extracts were prepared and subjected to immunoprecipitation with an anti-Flag antibody. The extracts (Input) and the immuno-precipitates (IP: Flag) were then subjected to western blotting analysis using antibodies as indicated. **h**, Schematic illustrating experiment design in Extended Data Fig. 8a. The weights of testes from 12-week-old *Ripk3^+/+^*, *Ripk3^-/-^*littermate and *Zbp1^-/-^* male mice (n=7 for each genotype) after heat shock treatment for 7 days. **i**, **j**, H&E staining of testis sections from 12-week-old *Ripk3^+/+^*, *Ripk3^-/-^* littermate and *Zbp1^-/-^* male mice after heat shock treatment for 7 days in (i). The thickness of seminiferous tubules was analyzed based on H&E staining and show in (j). scale bar, 100 μm. **k**, **l**, IHC staining of testis sections from 12-week-old *Ripk3^+/+^*, *Ripk3^-/-^* littermate and *Zbp1^-/-^* male mice with an anti-phospho-MLKL (p-MLKL) antibody after heat shock treatment for 7 days in (k). p-MLKL positive cells were counted in five fields per testis and quantified in (l). Scale bar, 100 μm. Data in (b, d and e) are mean ± SD of triplicate wells. Quantified data in (h, j and l) represent the mean ± s.e.m. \*\*\*\**P*<0.0001. *P* values were determined by two-sided unpaired Student’s *t* tests.

To dissect the molecular requirements for this process, we employed **IFN-β/HS treatment in L929 cells** lacking key necroptosis regulators. Genetic deletion of ***Ripk3*** or ***Mlkl*** abolished necroptosis, while ***Ripk1* knockout enhanced** IFN-β/HS-induced cell death (Fig. 2d), implicating **RIPK1 as a negative regulator** in this context. Consistently, **phospho-MLKL** levels were elevated in ***Ripk1*-deficient** L929 cells, while entirely absent in ***Ripk3*-null** cells following IFN-β/HS treatment (Extended Data Fig. 6d).

These findings were further supported in **MEF, GC-2spd, and 15P-1 cells**, all of which underwent **RIPK3-dependent necroptosis** in response to IFN-β/HS (Extended Data Fig. 6e–f). Collectively, these results demonstrate that **heat stress – induced necroptosis requires IFN-β – driven ZBP1 expression**, and proceeds through a **RIPK3 – MLKL – dependent mechanism**, with **RIPK1 acting as an inhibitory regulator** in this context.

### IFN-β/Heat Stress–Induced Necroptosis Requires Jak/STAT-Mediated ZBP1 Expression

To directly test the role of **ZBP1** in heat-induced necroptosis, we generated ***Zbp1*-**knockout L929, GC-2spd, and 15P-1 cells**. Upon** IFN-β and heat shock (IFN-β/HS) **treatment**, all *Zbp1*-deficient cell lines were completely **resistant to necroptotic cell death**, in contrast to wild-type controls (Fig. 2e, Extended Data Fig. 7a–b). ZBP1 protein was robustly induced in wild-type cells by IFN-β/HS, but absent in *Zbp1*-deficient counterparts (Fig. 2e, Extended Data Fig. 7a–b). Furthermore, **phospho-MLKL** was detected in IFN-β/HS-treated wild-type L929 cells, but **abolished in *Zbp1*-knockout cells**, confirming ZBP1 as essential for necroptosis induction (Fig. 2f).

Because **ZBP1 is an interferon-stimulated gene**, we next tested whether the **Jak/STAT signaling pathway** mediates its expression. Pharmacological inhibition of **Jak kinases** completely protected L929, GC-2spd, and 15P-1 cells from IFN-β/HS-induced cell death (Extended Data Fig. 7c–d). This protection was accompanied by a **loss of ZBP1 expression and phospho-MLKL**, indicating that **Jak/STAT signaling is required upstream of necroptosis in this context** (Extended Data Fig. 7e).

To confirm the sufficiency of ZBP1, we used **HeLa-RIPK3/TetOn-ZBP1 cells**, which stably express RIPK3 and allow doxycycline (DOX)-inducible ZBP1 expression. Upon **DOX/HS treatment**, these cells underwent robust **RIPK3-dependent necroptosis**, consistent with a direct role for ZBP1 in initiating the pathway (Extended Data Fig. 7f). We further confirmed that **ZBP1 physically interacts with RIPK3** following IFN-β/HS treatment using **co-immunoprecipitation**, demonstrating **ZBP1– RIPK3 complex formation** under necroptotic conditions (Fig. 2g). To dissect the structural requirements for ZBP1 function^21^, we introduced various **ZBP1 domain mutants** into ***Zbp1⁻/⁻* L929 cells**: full-length ZBP1, **ZBP1(ΔZα1/2)** (lacking both Zα domains), **ZBP1(ΔC)** (lacking the C-terminal domain), and **ZBP1mut** (bearing point mutations in both RHIM domains) (Extended Data Fig. 7g). Re-expression of wild-type ZBP1 or ZBP1(ΔC) restored necroptosis, while **ZBP1(ΔZα1/2)** and **ZBP1mut** failed to do so—even though all constructs were expressed at similar levels (Extended Data Fig. 7g–h). These results indicate that both the **Zα domains** and **RHIM motifs** are required for **ZBP1-mediated activation of RIPK3 and necroptosis** in response to heat stress. Finally, unlike the cell lines requiring IFN-β priming, **ZBP1 is constitutively expressed in human and mouse testicular tissue** (Extended Data Fig. 7i–j). This **basal ZBP1 expression** allows for **direct activation of ZBP1 – RIPK3 – MLKL – mediated necroptosis** *in vivo*, helping to explain the susceptibility of testicular cells to heat-induced cell death and azoospermia.

### *Zbp1* or *Ripk3* Deficiency Protects Against Heat-Induced Testicular Damage *In Vivo*

To assess whether **heat stress triggers ZBP1–RIPK3–dependent necroptosis *in vivo***, we established a mouse model in which the **scrotum was immersed in 43°C water** for 20 minutes **every other day over five days**^22–24^. Mice were euthanized on day 7 following the final exposure, and their **body weights, testicular weights, and histological features** were examined (Extended Data Fig. 8a–b). In **wild-type mice**, heat exposure led to **significant testicular atrophy**, as indicated by reduced testis weight. In contrast, both ***Ripk3*-knockout** and ***Zbp1*-knockout** mice were **largely protected from weight loss** (Fig. 2h, Extended Data Fig. 8c). Histological analysis revealed **severe cell depletion** and near-empty **seminiferous tubules** in heat-treated wild-type testes, whereas **tubules in knockout mice remained intact**, with lumens containing abundant germ cells and spermatozoa—resembling untreated controls (Fig. 2i, Extended Data Fig. 8d). Heat-induced **thinning of seminiferous tubule walls** was also significantly attenuated in ***Ripk3*- and *Zbp1*-deficient mice** (Fig. 2j). Furthermore, **phospho-MLKL** was robustly detected in the **seminiferous epithelium** of wild-type testes post-heat stress but was **undetectable** in the same regions of ***Ripk3*- and *Zbp1*-deficient** testes (Fig. 2k–l). These findings demonstrate that **heat stress activates** necroptosis in testicular tissue via a ZBP1–RIPK3–MLKL axis**, and that** genetic **ablation of *Zbp1* or *Ripk3* is sufficient to prevent heat-induced testicular damage and germ cell loss**, supporting a central role for this pathway in heat-related azoospermia.

### Stress Granule Formation Is Required for Heat-Induced Necroptosis

**Stress granules (SGs)** are membrane-less, phase-separated cytoplasmic assemblies of ribonucleoproteins that form in response to acute stressors such as **heat shock, viral infection, tumorigenesis**, and **neurodegeneration**^25,26^. Key RNA-binding proteins, including **TIA-1** and **G3BP1**, are essential for SG nucleation^27,28^. Although SGs have been proposed to promote **cell survival**^29^, recent work has shown that certain cytotoxic insults, such as **sodium arsenite**, can trigger **ZBP1–RIPK3–dependent necroptosis** via recruitment of RIPK3 to SGs^30^.

To investigate whether SGs are similarly required for **heat-induced necroptosis**, we treated **L929 cells** with **IFN-β and heat shock (IFN-β/HS)** following *G3bp1*, *G3bp2*, or *G3bp1/2* double-knockout (DKO). Western blot analysis showed that *G3bp1/2*-DKO cells were completely resistant to IFN-β/HS-induced necroptosis and failed to activate **phospho-MLKL** (Fig. 3a–b, Extended Data Fig. 9a–b).

**Fig. 3.**
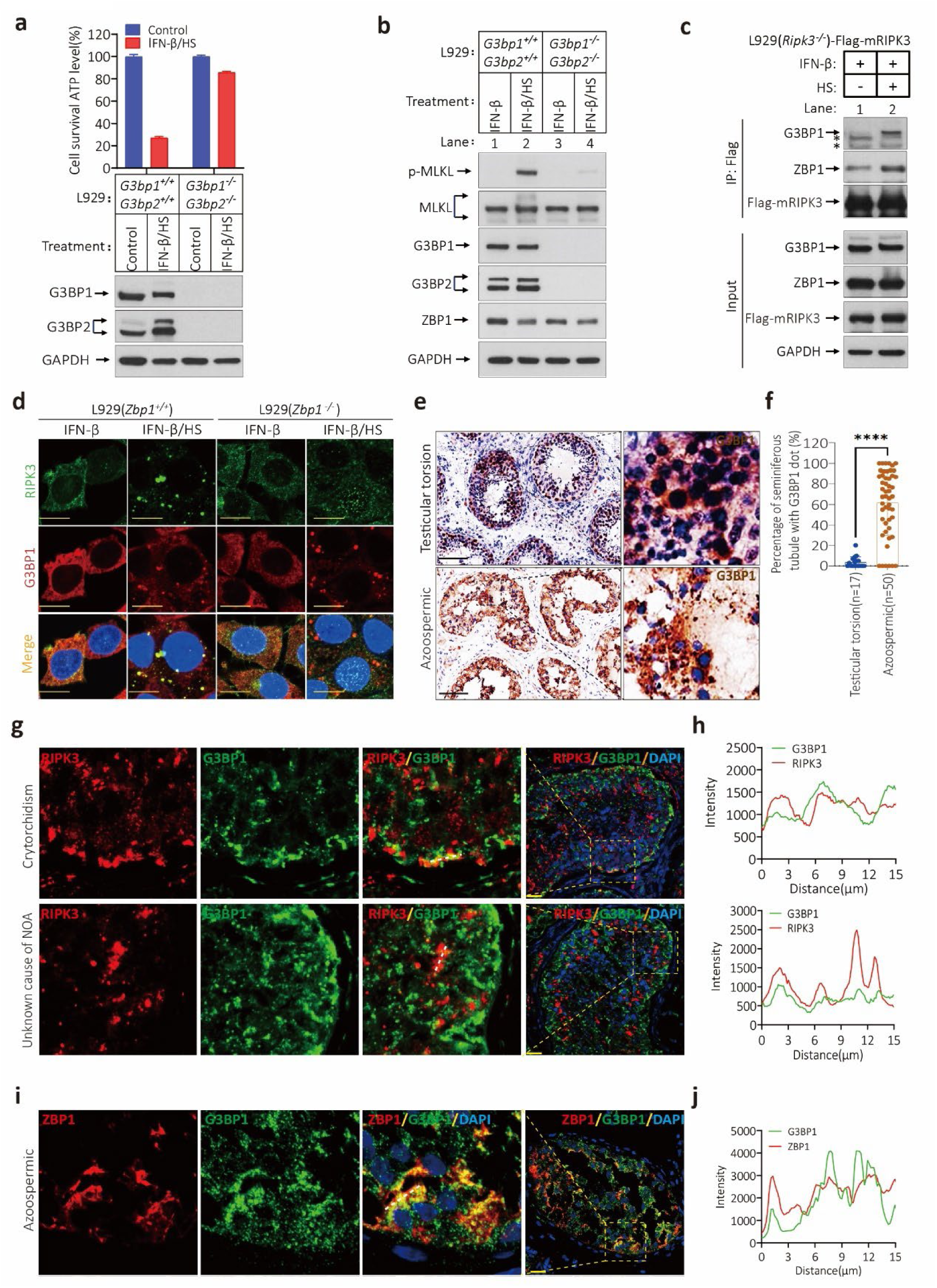
Stress granule is required for heat shock-induced necroptosis. **a**, **b**, Cultured L929 cells with wild type or *G3bp1* and *G3bp2* double gene knocked out were treated with IFN-β and HS for 18 hours and 1.5 hours as described in Extended Data Fig. 6a, 2 hours after heat shock cell viability as measured by Cell Titer-Glo (a). The levels of p-MLKL, MLKL, ZBP1, G3BP1, and G3BP2 were analyzed by immunoblotting in (b), GAPDH was used as loading control. Data in (A) are mean ± SD of triplicate wells. **c**, Cultured L929(*Ripk3^-/-^*)-HA-3×Flag-mRIPK3 cells were treated with IFNβ/HS as indicated. The cell extracts were prepared and subjected to immunoprecipitation with an anti-Flag antibody. The extracts (Input) and the immuno-precipitates (IP: Flag) were then subjected to western blotting analysis using antibodies as indicated. The asterisk(*) indicates non-specific bands. **d**, Cultured L929(*Zbp1^+/+^*) and L929(*Zbp1^-/-^*) cells were treated with the indicated stimuli for 18 hours (IFN-β) and 0.5 hours (HS). The RIPK3 and G3BP1 were detected by immunofluorescence. Scale bares, 10 μm. **e**, **f,** IHC analysis of human testicular torsion and NOA testis sections with G3BP1 antibody in (e). The number of seminiferous tubules with positive G3BP1 dot signal were counted based on IHC staining and quantification in (f). Scale bar, 100 μm. Data represent the mean ± s.e.m. \*\*\*\**P*<0.0001. *P* values were determined by two-sided unpaired Student’s *t* tests. **g**, **h**, Immunofluorescence analysis of NOA testis sections (unknown cause of NOA, n=5; crytorchidism, n=5) with antibodies against G3BP1(green) and RIPK3(red) in (g). Scale bar, 50 μm. Profiling of representative white dotted line traces the intensities of RIPK3 and G3BP1 signal based on (g) and analyzed in (h). **i**, **j**, Immunofluorescence analysis of NOA testis sections (unknown cause of NOA, n=5; crytorchidism, n=5) with antibodies against G3BP1(green) and ZBP1(red) in (i). Scale bar, 50 μm. Profiling of representative white dotted line traces the intensities of ZBP1 and G3BP1 signal based on (i) and analyzed in (j).

**Co-immunoprecipitation experiments** revealed that **RIPK3** was recruited to SGs and formed a complex with **ZBP1** and **G3BP1** following IFN-β/HS treatment (Fig. 3c), implicating a functional interaction. To investigate how RIPK3 is targeted to SGs, we examined the subcellular localization of RIPK3 and G3BP1 in **L929, GC-2spd**, and **15P-1 cells**, with or without *Zbp1*, before and after IFN-β/HS exposure. In untreated cells, **RIPK3 was diffusely cytoplasmic**, but after treatment, it became **punctate and filamentous**, co-localizing with G3BP1 (Fig. 3d, Extended Data Fig. 9c). This redistribution was **abolished in *Zbp1*-deficient cells**, indicating that **ZBP1 is required for RIPK3 recruitment to stress granules**.

To assess whether SG formation occurs ***in vivo***, we performed **IHC staining** of **NOA patient testes** using a **G3BP1 antibody**. Punctate G3BP1 signals were prominently detected in the seminiferous tubules of most NOA samples but were mostly **absent** in testicular torsion controls (Fig. 3e–f). Importantly, these **G3BP1 puncta co-localized with ZBP1 and RIPK3** within the same testicular cells, including those from patients without cryptorchidism (Fig. 3g – j). Similarly, **heat-treated mouse testes** exhibited robust G3BP1 puncta, which were **nearly absent in untreated controls** (Extended Data Fig. 9d–f). Collectively, these results demonstrate that **stress granules are essential platforms for ZBP1 – RIPK3 complex formation and necroptosis activation**, both ***in vitro* and *in vivo***, and that this mechanism underlies testicular degeneration in **NOA**.

### eIF2α Kinases Are Required for Stress-Induced Necroptosis

The **integrated stress response (ISR)** is a cytoprotective signaling cascade triggered by phosphorylation of **eukaryotic initiation factor 2 alpha (eIF2α)** at **Ser51**, a key regulatory event that inhibits translation during stress^31,32^. Four eIF2α kinases mediate ISR activation in response to distinct stimuli: **HRI** (heme depletion), **PKR** (viral infection), **PERK** (ER stress), and **GCN2** (amino acid deprivation)^32^. Deletion of all four kinases abrogates ISR signaling entirely^33^.

To examine whether **eIF2α kinases are required for necroptosis**, we generated **quadruple-knockout L929 cells** (lacking HRI, PKR, PERK, and GCN2; Extended Data Fig. 10a). Upon **IFN-β/heat shock (HS) treatment**, **quadruple-knockout cells were resistant to necroptosis**, and **phospho-MLKL** was not detected (Fig. 4a–b). Stepwise deletion revealed that **PKR and HRI deficiency partially reduced** necroptosis, while **GCN2 knockout** — either alone or in combination — **almost completely abolished** cell death. In contrast, **PERK knockout had no additional protective effect** (Extended Data Fig. 10b). Consistently, **phospho-MLKL, phospho-eIF2α, and G3BP1 puncta** were markedly reduced in GCN2-deficient cells but not in PKR/HRI/PERK-deficient cells (Fig. 4b–c, Extended Data Fig. 10c). We next tested whether other ISR-activating stressors could trigger ZBP1 – RIPK3 – dependent necroptosis. Exposure to **arsenite**, **halofuginone** (GCN2-dependent), and **8-azaadenosine** (PKR-dependent) all induced necroptosis in L929 cells, while **tunicamycin** (PERK-dependent) did not (Extended Data Fig. 11a–d). These results suggest that **GCN2, PKR, and HRI — but not PERK — mediate necroptosis induction** under stress. To explore whether ISR is similarly activated *in vivo*, we performed IHC staining of **testis samples from NOA patients**. **Phosphorylated GCN2 (p-GCN2)** was detected in 90% of NOA samples (45/50), while **phospho-PKR** was detected in 46% (23/50) (Fig. 4j, k). Both signals were nearly absent in **testicular torsion controls** (Fig. 4d–e, g–h). **Phospho-GCN2 and phospho-PKR co-localized with phospho-MLKL** in the same seminiferous tubule cells, supporting their role in necroptosis (Fig. 4f, i). Detection of **phospho-PERK** was minimal in both NOA and control samples (Extended Data Fig. 11e–f) and **HRI phosphorylation could not be evaluated** due to a lack of specific antibodies. These findings demonstrate that activation of the **ISR**, particularly through **GCN2**, is essential for **stress granule formation** and **ZBP1–RIPK3-mediated necroptosis**. The presence of activated eIF2 α kinases in NOA testis supports a model in which **chronic or acute stress triggers ISR-driven necroptosis**, contributing to **testicular degeneration and infertility**.

**Fig. 4.**
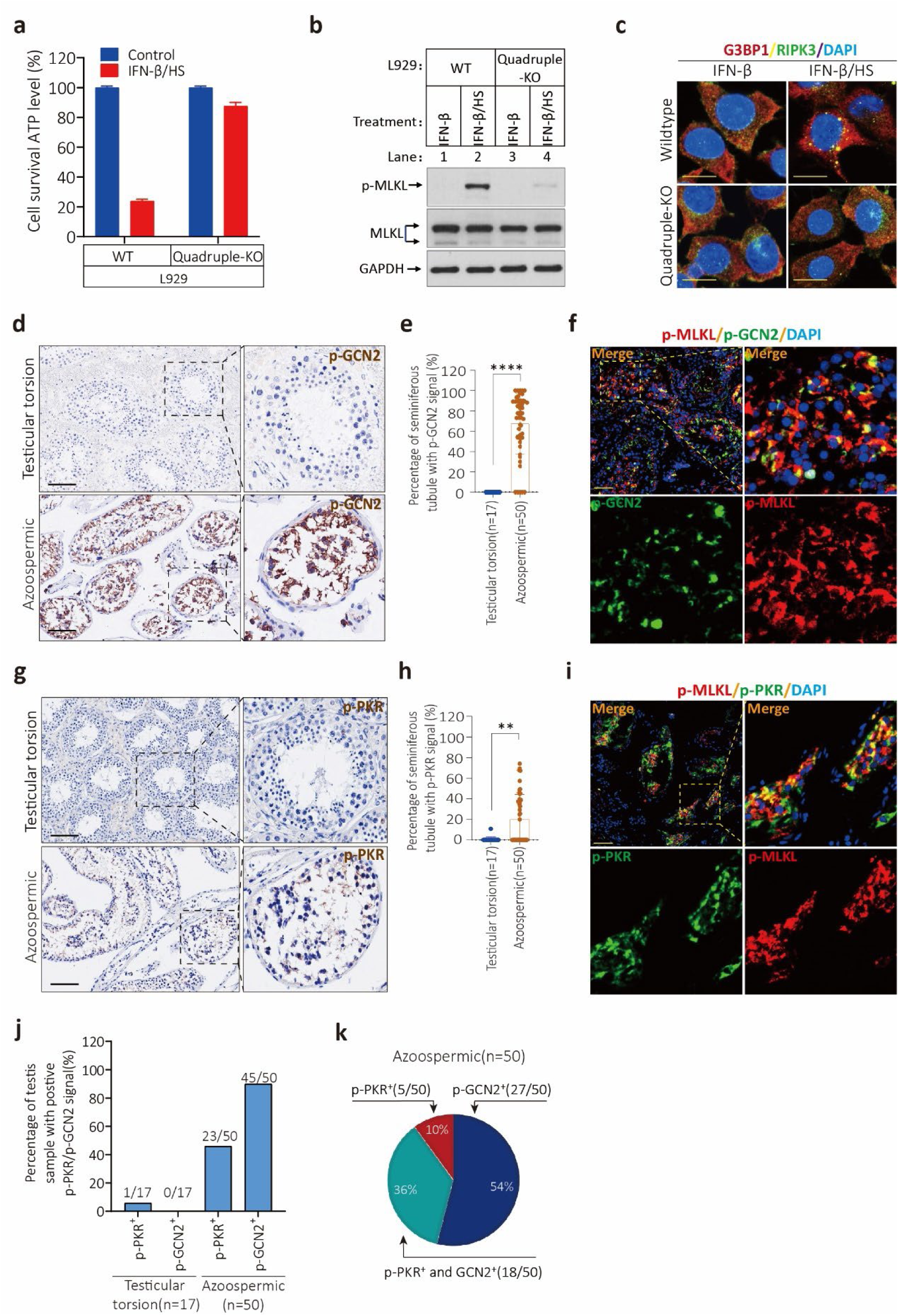
Stress kinases induce ZBP1 and RIPK3-dependent necroptosis in non-obstructive azoospermia. **a**, **b**, Cultured L929 cells with wildtype or *Pkr*, *Hri*, *Perk* and *Gcn2* four gene knocked out (quadruple-KO) were treated with IFN-β and HS for 18 hours and 1.5 hours as described in Extended Data Fig.6a, 2 hours after heat shock cell viability as measured by Cell Titer-Glo (a). The levels of p-MLKL and MLKL were analyzed by immunoblotting in (b), GAPDH was used as loading control. Data in (a) are mean ± SD of triplicate wells. **c**, Cultured L929(wildtype) and L929(quadruple-KO) cells were treated with the indicated stimuli for 18 hours (IFN-β) and 0.5 hours (HS). The RIPK3 and G3BP1 were detected by immunofluorescence. Scale bares, 10 μm. **d**, **e**, IHC analysis of human testicular torsion and NOA testis sections with p-GCN2 antibody in (d). The number of seminiferous tubules with positive p-GCN2 signal were counted based on IHC staining and quantification in (e). Scale bar, 100 μm. Data represent the mean ± s.e.m. \*\*\*\**P*<0.0001. *P* values were determined by two-sided unpaired Student’s *t* tests. **f**, Immunofluorescence analysis of NOA testis sections(n=5) with antibodies against p-MLKL (red) and p-GCN2(green). Scale bar, 50 μm. **g**, **h**, IHC analysis of human testicular torsion and NOA testis sections with p-PKR antibody in (g). The number of seminiferous tubules with positive p-PKR signal were counted based on IHC staining and quantification in (h). Scale bar, 100 μm. Data represent the mean ± s.e.m. \*\**P* < 0.01. *P* values were determined by two-sided unpaired Student’s *t* tests. **i**, Immunofluorescence analysis of NOA testis sections(n=5) with antibodies against p-MLKL (red) and p-PKR (green). Scale bar, 50 μm. **j**, **k**, Analysis of p-GCN2 and p-PKR positive testis sections from the testicular torsion(n=17) and NOA(n=50) testes in (j). Analysis of NOA testis sections with single positive p-GCN2, single positive p-PKR and double positive p-GCN2 and p-PKR in (k).

### Stress Markers and Phospho-MLKL Are Elevated in Aging Human Testes

To investigate whether **ZBP1 – RIPK3 – dependent necroptosis** occurs during physiological testis aging, we analyzed **testicular tissue from 30 prostate cancer patients** (age range: 59–89 years; mean: 73; Supplementary Table 1). Testicular torsion samples served as controls. Histological analysis revealed that **aging testes had significantly lower Johnsen Scores** (mean score = 7.4), indicating **impaired spermatogenesis** compared to controls (Fig. 5a– b). Immunohistochemistry (IHC) using a **human phospho-MLKL antibody** revealed robust staining in the seminiferous tubules of **29/30 (96.7%) aging testis samples**, whereas **no signal** was detected in torsion controls (0%, n = 17) (Fig. 5c–d). We next asked whether the **ISR–stress granule–necroptosis pathway** was also activated in aging testes. IHC for **G3BP1**, **phospho-GCN2**, **phospho-PKR**, and **phospho-PERK** were performed. **Punctate G3BP1 staining**, indicative of stress granule formation, was detected in most aging samples but was **absent or minimal in torsion controls** (Fig. 5e – f). Similarly, **phospho-GCN2** was present in **96.7% (29/30)** of aging samples, and **phospho-PKR** in **23% (7/30)**. In contrast, **phospho-PERK** was not detected in either group (Fig. 5g–h, j–k, Extended Data Fig. 11g–j). Co-localization analysis revealed that **phospho-GCN2 and phospho-PKR overlapped with phospho-MLKL** in the same seminiferous tubule cells (Fig. 5i, Extended Data Fig. 11k), suggesting functional integration of these stress pathways. Of the 30 aging samples, **29** exhibited p-GCN2, **7** exhibited p-PKR, and **6** were positive for both (Fig. 5j–k). Together, these findings demonstrate that **eIF2α kinase activation and stress granule formation occur in the aging human testis**, leading to **ZBP1–RIPK3–MLKL–mediated necroptosis**. These results suggest that testis aging shares a common molecular pathway with NOA, driven by **chronic stress signaling and regulated cell death** (Extended Data Fig. 12).

**Fig. 5.**
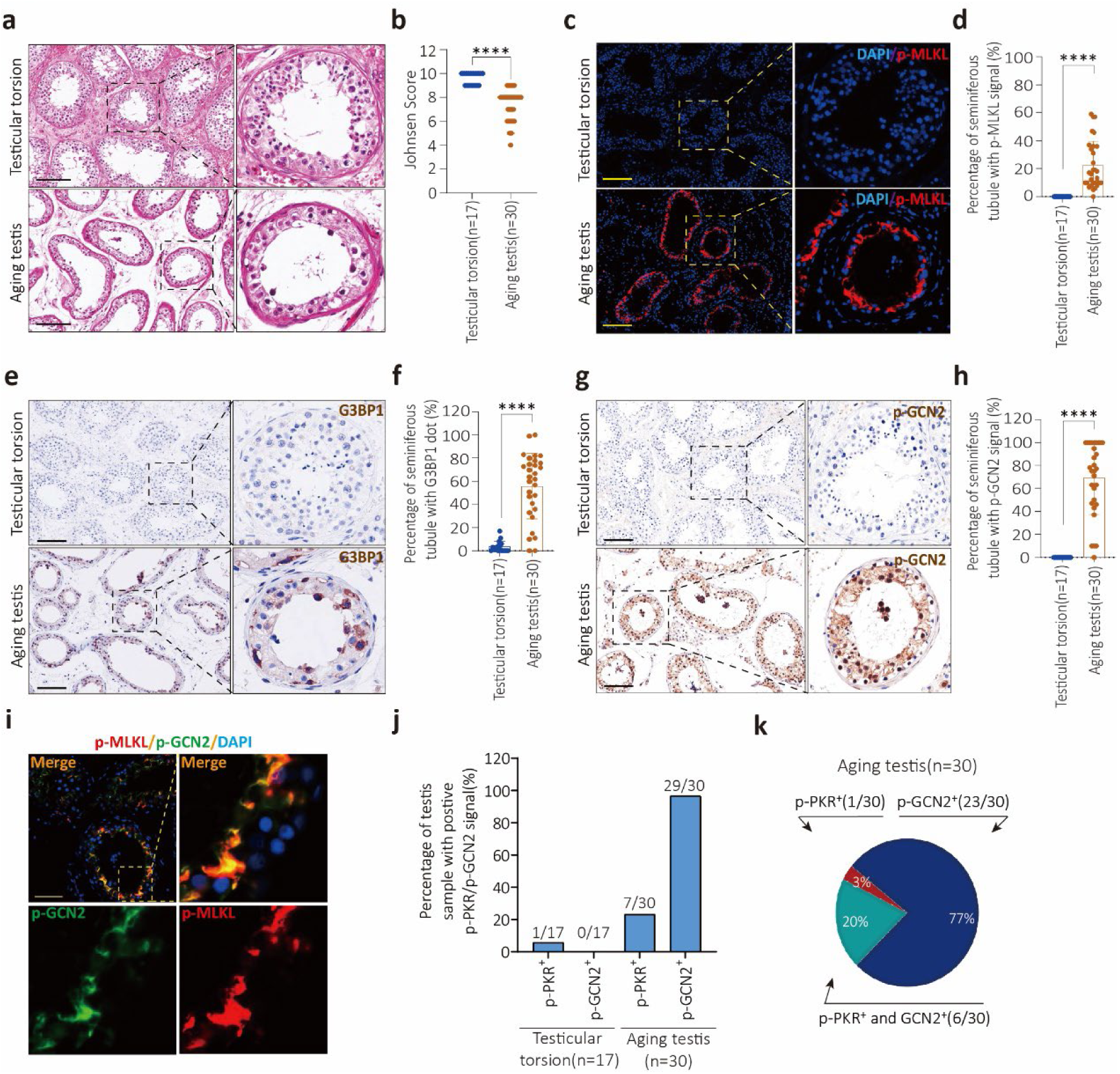
p-MLKL signals were associated with stress biomarkers in aging testes. **a**, **b**, H&E staining of testis sections from human testicular torsion(n=17) and prostate cancer patients (aging testis, n=30) in (a). Johnsen Score evaluation of testicular torsion and prostate cancer patients based on (a) and show in (b). Scale bar, 100 μm. **c**, **d**, Immunofluorescence analysis of human testicular torsion and aging testis sections with p-MLKL antibody in (c). The number of seminiferous tubules with positive p-MLKL signal were counted based on immunofluorescence staining and quantification in (d). Scale bar, 100 μm. **e**, **f**, IHC analysis of human testicular torsion and aging testis sections with G3BP1 antibody in (e). The number of seminiferous tubules with positive G3BP1 dot signal were counted based on IHC staining and quantification in (f). Scale bar, 100 μm. Data represent the mean ± s.e.m. \*\*\*\**P*<0.0001. *P* values were determined by two-sided unpaired Student’s *t* tests. **g**, **h**, IHC analysis of human testicular torsion and aging testis sections with p-GCN2 antibody in (g). The number of seminiferous tubules with positive p-GCN2 signal were counted based on IHC staining and quantification in (h). Scale bar, 100 μm. Data represent the mean ± s.e.m. \*\*\*\**P*<0.0001. *P* values were determined by two-sided unpaired Student’s *t* tests. **i**, Immunofluorescence analysis of aging testis sections(n=5) with antibodies against p-MLKL (red) and p-GCN2(green). Scale bar, 50 μm. **j**, **k**, Analysis of p-GCN2 and p-PKR positive testis sections from the testicular torsion(n=17) and aging testes(n=30) in (j). Analysis of aging testis sections with single positive p-GCN2, single positive p-PKR and double positive p-GCN2 and p-PKR in (k).

## Discussion

Our findings identify **ZBP1 – RIPK3 – dependent necroptosis** as a central mechanism underlying **non-obstructive azoospermia (NOA)**, one of the most severe forms of male infertility. The necroptosis marker **phospho-MLKL** was detected in **100% of NOA patient testis samples** but not in any controls, indicating that this pathway is consistently activated across a clinically and etiologically heterogeneous patient population. Phospho-MLKL localized specifically to **spermatogonia and Sertoli cells**—two cell types essential for spermatogenesis—strongly implicating necroptotic cell death as a direct cause of germ cell loss in NOA.

Mechanistic studies in cell lines and mouse models revealed that **necrotic cell death in the testis is triggered by stress granules**, which act as scaffolds to recruit and activate **ZBP1**, leading to downstream activation of **RIPK3 and MLKL**. SG formation itself is initiated by **eIF2α kinase signaling**, particularly through **GCN2** and **PKR**, which respond to diverse stressors including **heat shock**, **oxidative stress**, and **amino acid deprivation**. Markers of this stress axis— including **phospho-GCN2**, **phospho-PKR**, **punctate G3BP1**, **RIPK3**, and **phospho-MLKL**—were consistently detected and **co-localized** within the same testicular cell types in NOA patients.

Despite the clinical heterogeneity of NOA, the uniform activation of necroptosis across all patients suggests the presence of a **common molecular effector pathway**. Our results support a model in which **diverse environmental and endogenous stress signals** converge on **eIF2α kinase–driven stress granule assembly**, which in turn activates the **ZBP1–RIPK3–MLKL** cascade. ZBP1 is likely recruited to SGs via its **Z α domain**, forming necrosomes with RIPK3 and initiating cell death. Notably, multiple eIF2α kinases were activated in several patient samples, with **GCN2** being the most consistently observed. This raises the possibility that **NOA may reflect accumulated responses to a variety of environmental or intrinsic insults**, such as **hyperthermia**, **nutrient stress**, **viral infections**, or **endogenous retro-element activation**. While we could not assess **HRI activity** in patient samples due to technical limitations, our cell-based assays indicate it may also participate in this pathway.

Importantly, the necroptosis pathway characterized here in NOA mirrors that observed in **physiological testicular aging**. Phospho-MLKL and upstream stress markers were also detected in aging testes from older men, suggesting that **chronic, low-level activation of the SG– ZBP1 – RIPK3 pathway** may contribute to **age-associated testicular decline**. These findings position NOA as a **model of premature testicular aging** and offer a mechanistic framework for future studies of male reproductive aging. Together, our study uncovers a **stress-induced, necroptotic cell death pathway in the testis**, centered on **ZBP1 recruitment to stress granules**, and **activated by eIF2α kinases**. This axis offers a unifying explanation for both **NOA and natural aging**, and highlights potential molecular targets for preserving male fertility under environmental or physiological stress.

## Acknowledgments and Funding

This work was supported by institutional grants from the Chinese Ministry of Science and Technology and Beijing Municipal Commission of Science and Technology. The funders had no role in study design, data collection and interpretation, or the decision to submit the work for publication.

## Author contributions

D.L, L.T and X.W. conceived the project, supervised the research, and wrote the manuscript; D.L. and H.L designed the experiments; H.L., D.L., J.C., B.D., K.J., T.X., and H.H. performed the experiments; W.F. and L.T. commented on and edited the manuscript; H.L. and D.L. analyzed the data and made the figures.

## Data and materials availability

All data are available in the main text or supplementary materials. Correspondence and requests for materials should be addressed to D.L, L.T and X.W.

## Methods and Extended Data Figs

### Methods Mice

The *Ripk3^-/-^* and *Zbp1^-/-^*mice (C57BL/6NCrl strain) were kept in our lab. The primers used for genotyping are listed below.

Mouse *Ripk3*-KO-F: CAGTGGGACTTCGTGTCCG

Mouse *Ripk3*-KO-R: CAAGCTGTGTAGGTAGCACATC

Mouse *Zbp1*-WT-F: AGAGTTGGGGGTTCCTACCT

Mouse *Zbp1*-WT-R: TGAGGGTTTTCTTGGGCACT

Mouse *Zbp1*-KO-F: GTGGCTGAAGCAGGAGGATT

Mouse *Zbp1*-KO-R: ATTGGTAGCCCTTGTGAGGC

### Animal husbandry

Mice were group-housed in a 12 hours light/dark (light between 08:00 and 20:00) in a temperature-controlled room (21.1 ± 1 °C) at the Sironax with free access to water. The ages of mice are indicated in the figure, figure legends, or methods. All animal experiments were conducted following the Ministry of Health national guidelines for the housing and care of laboratory animals and were performed in accordance with institutional regulations after review and approval by the Institutional Animal Care and Use Committee at the Sironax, Beijing.

### Heatstroke model

Adult male C57BL/6 mice (12 weeks) weighted 25∼30g were anesthetized by injecting 1.25% Avertin (2,2,2-tribromoethanol, M2920, Aibei Biotechnology, Nanjing, China) intraperitoneally at concentrations of 0.12 ml/10g. Then, the lower parts of the body (hind legs, tail, and scrotum) were submerged in a thermostatically controlled water bath at 43°C or 37°C for 20 min, every other day for a total of three times (Extended Data Fig. 8a). Immediately after the heat stress exposure, mice were subsequently moved back to their original cages with an environmental temperature at 21.1 ± 1 °C and free access to food and water. 7 days after the final heatstroke, mice were sacrificed and perfused with PBS. Testes were weighed and fixed in Bouin’s solution for histological and immunohistochemical assays.

### Human tissues

The research involving human tissue samples were dissected from human testicular torsion, azoospermia and prostate cancer patients (n=17, 13-30 years, testicular torsion patients; n=50, 18-57 years, azoospermia patients; n=30, 59-89 years, prostate cancer patients) were provided from Beijing Chao-Yang Hospital in China and snap-frozen in liquid nitrogen and stored at -80°C. Tissues were cut into appropriately-sized pieces and placed in Bouin’s solution for preservation. After several days of Bouin’s solution fixation at room temperature, tissue fragments were transferred to 70% ethanol and stored at 4°C.

The medical ethics committee of the Beijing Chao-Yang Hospital and National Institute of Biological Sciences, Beijing, China approved the study (2022-KE-491).

### Antibodies and reagents

Antibodies used in this study were anti-GAPDH-HRP (M171-1, MBL; WB, 1:5000), anti-Flag (F1804, Sigma-Aldrich; WB, 1:5000), anti-Human-p-RIPK1 (#44590S, Cell Signaling; IHC, 1:100), Cleaved Caspae3 (#9661L, Cell Signaling; IHC, 1:100), anti-RIPK3 (#2283, ProSci; WB, 1:1000; IHC, 1:100), anti-Mouse-RIPK3 (#95702, Cell Signaling; WB, 1:1000), anti-MLKL (ab255747; abcam; WB, 1:1000), anti-Mouse-p-MLKL (ab196436; WB, 1:1000; IHC, 1:100), anti-Human-p-MLKL (ab187091; WB, 1:1000; IHC, 1:100), anti-Human/Mouse-ZBP1 (AG-20B-0010-C100, AdipoGen; WB, 1:1000; IHC, 1:100), anti-G3BP1 (13057-2-AP, proteintech; WB, 1:1000; IHC, 1:100), anti-G3BP2 (PA5-53797, Invitrogen; WB, 1:1000), anti-PKR (ab184257, abcam; WB, 1:1000), anti-PKR (Abways, CY5665; WB, 1:1000), anti-PERK (377400, Santa Cruz Biotechnology; WB, 1:1000), anti-GCN2 (3302, Cell Signaling; WB, 1:1000), anti-PIWIL4 (PA5-31448, thermo; IHC, 1:500), anti-Sox9 (ab185966, abcam; IHC, 1:100), DDX4 (ab13840, abcam; IHC, 1:100), anti-p-GCN2(T899) (ThermoFisher, PA5-105886; WB, 1:1000; IHC, 1:500), anti-p-PKR (Abways, CY5271; WB, 1:1000; IHC, 1:100), anti-p-PERK(T981) (CUSABIO, CSB-PA072558; IHC, 1:100), Donkey anti-Mouse, Alexa Fluor 488 (Thermo Fisher, A-21202), Donkey anti-Mouse, Alexa Fluor 555 (Thermo Fisher, A-31570), Donkey anti-Rabbit, Alexa Fluor 488 (Thermo Fisher, A-21206), and Donkey anti-Rabbit, Alexa Fluor 555 (Thermo Fisher, A-31572). Reagents used in this study were INF-β (Sino Biological, 50708-MCCH), Baricitinib (JAK inhibitor, MedChemExpress, HY-15315), DOX (Sigma, WXBC9363V), NaAsO_2_ (Sodium arsenite, Innochem, A25410), Halofuginone (MedChemExpress, S8144), 8-Azaadenosine (MedChemExpress, HY-115686), Tunicamycin (MedChemExpress, S7894) and Lambda Protein Phosphatase (Cat: P0753S, 400,000 U/mL). Phoshorylated MLKL peptides (GYQVKLAGFELRKTQpTpSMSLGTTREKTDRVKS) were synthesized by GenScript.

### Constructs

psPAX2 and pMD2.G construct were kept in our lab. Full-length mouse ZBP1 and truncated ZBP1(lacked two Zα domain or C-terminal domain (315-411)) were subcloned into the pWPI (GFP-tagged) vector to generate pWPI-mZBP1 (WT, ΔZα and ΔC) construct. Using Quickchange Site-Directed Mutagenesis Kit to generate pWPI-mZBP1^mut^ (two points mutation within two RHIM domain) construct.

The gRNAs for targeting mouse *Zbp1*, *G3bp1/2*, *Pkr*, *Perk*, *Gcn2* and *Hri* were designed and were cloned into the gRNA-Cas9 expression plasmid pX458-GFP to generate pX458-GFP-ZBP1/G3BP1/G3BP2/PKR/PERK/GCN2/HRI construct. The sequences used for gRNAs targeting are listed below.

Mouse ZBP1-gRNA: GAAGATCTACCACTCACGTC

Mouse G3BP1-gRNA: ATGTTCACAACGACATCTTC

Mouse G3BP2-gRNA: AAGCTCCCGAGTATTTGCAC

Mouse HRI-gRNA: TCGAAGCACAAACGTCACGC

Mouse PKR-gRNA: TTGTTCGTTGGTAACTACAT

Mouse PERK-gRNA: CTCGAATCTTCCTACAAGTT

Mouse GCN2-gRNA: AAAGCCCGGACATACTCCTC

### Cells

All cells were cultured at 37°C with 5% CO_2_. All cell lines were cultured as follows: HEK293T (293T) were obtained from ATCC and cultured in DMEM (Hyclone). Mouse embryonic fibroblasts (MEF), MEF(*Ripk3^-/-^*) and HeLa-RIPK3/Teton-ZBP1 cells were cultured in DMEM. L929, L929(*Ripk1^-/-^*), L929(R*ipk3^-/-^*), L929(*Mlkl^-/-^*), L929(*Zbp1^-/-^*), L929(*Pkr^-/-^*), L929(*Pkr^-/-^Hri^-/-^*), L929(*Pkr^-/-^Hri^-/-^Perk^-/-^*) and L929(*Pkr^-/-^ Hri^-/-^Perk^-/-^Gcn2^-/-^*) were cultured in DMEM. GC-2spd(ts) and 15P-1 cells were obtained from ATCC and cultured in DMEM. GC-2spd(*Zbp1^-/-^*) and 15P-1(*Zbp1^-/-^*) were cultured in DMEM. L929(*Zbp1^-/-^*) cells were infected with virus encoding ZBP1 (WT, RHIM^mut^, ΔZα and ΔC) to establish the L929(*Zbp1^-/-^*)-ZBP1, L929(*Zbp1^-/-^*)- ZBP1(RHIM^mut^), L929(*Zbp1^-/-^*)-ZBP1(ΔZα) and L929(*Zbp1^-/-^*)-ZBP1(ΔC) cell lines. L929(*Ripk3^-/-^*) cells were infected with virus encoding HA-3×Flag-mRIPK3 and GFP-positive live cells were sorted to establish the L929(*Ripk3^-/-^*)-HA-3×Flag-mRIPK3 cell lines. All media were supplemented with 10% FBS (Thermo Fisher) and 100 units/ml penicillin/ streptomycin (Thermo Fisher). MA-10 obtained from ATCC and cultured in DMEM:F12 (Hyclone, additional 20 mM HEPES, horse serum to a final concentration of 15%).

### Cell survival assay

Cell survival assay was performed using Cell Titer-Glo Luminescent Cell Viability Assay kit. A Cell Titer-Glo assay (Promega, G7570) was performed according to the manufacturer’s instructions. Luminescence was recorded with a Tecan GENios Pro plate reader.

### CRISPR/Cas9 knockout cells

10 μg of pX458-GFP-ZBP1/G3BP1/G3BP2/PKR/PERK/GCN2/HRI plasmid was transfected into 1×10^5^ L929 cells using the Neon™ transfection system (Invitrogen™, MPK5000) by following the manufacturer’s instructions. 3 days after the transfection, GFP-positive live cells were sorted into single clones by using a BD FACSArial cell sorter. The single clones were cultured into 96-well plates for another 10-14 days or longer, depending upon the cell growth rate. The anti-ZBP1/G3BP1/G3BP2/PKR/PERK/GCN2 immunoblotting was used to screen for the L929(*Zbp1^-/-^*), L929(*Pkr^-/-^*), L929(*Pkr^-/-^Hri^-/-^*), L929(*Pkr^-/-^Hri^-/-^Perk^-/-^*) and L929(*Pkr^-/-^Hri^-/-^Perk^-/-^Gcn2^-/-^*) clones. Genome type of the knockout cells was determined by DNA sequencing.

### Cell stress exposure

Cultured L929, L929(*Pkr^-/-^*), L929(*Pkr^-/-^Hri^-/-^*), L929(*Pkr^-/-^Hri^-/-^Perk^-/-^*), L929(*Pkr^-/-^ Hri^-/-^Perk^-/-^Gcn2^-/-^*), L929(*G3bp1^-/-^G3bp2^-/-^*), L929(*Zbp1^-/-^*), L929(*Ripk3^-/-^*), L929(*Mlkl^- /-^*), MEF, MEF(*Ripk3^-/-^*), HeLa-RIPK3/Teton-ZBP1, GC-2spd, 15P-1, GC-2spd(*Zbp1^-/-^*), 15P-1(*Zbp1^-/-^*) and MA-10 cells were pretreated with DMSO, Dox or recombinant mouse INF-β (50708-MCCH, Sino Biological, Beijing, China) at 10 ng/ml for 18 hours before were trypsinized and resuspended with the complete medium. Cells were placed in a water bath with a temperature at 43°C or 37°C for the indicated times, and then incubated at 37°C and humidified 5% CO_2_ for the indicated time periods. Finally, cell lysates and supernatants were collected at the indicated time points after heat stress for ATP activity, western-blot, immunoprecipitation, or immunofluorescence assay.

Cultured L929, L929(*Pkr^-/-^Hri^-/-^Perk^-/-^Gcn2^-/-^*), L929(*G3bp1^-/-^G3bp2^-/-^*), L929(*Zbp1^-/-^*), L929(*Ripk3^-/-^*) and L929(*Mlkl^-/-^*) cells were treated with NaAsO_2_(30 μM) and NaAsO_2_(30 μM)+IFN-β(10 ng/ml) for 18 hours; Halofuginone(500 nM) and Halofuginone(500 nM)+IFN-β(10 ng/ml) for 42 hours; 8-Azaadenosine (20 μM) and 8-Azaadenosine(20 μM)+IFN-β(10 ng/ml) for 46 hours; Tunicamycin (2.5 μM) and Tunicamycin(2.5 μM)+IFN-β(10 ng/ml) for 30 hours. The intracellular ATP levels were measured by Cell Titer-Glo.

### Western blotting

Cell pellet samples were collected and re-suspended in lysis buffer (100 mM Tris-HCl, pH 7.4, 100 mM NaCl, 10% glycerol, 1% Triton X-100, 2 mM EDTA, Roche complete protease inhibitor set, and Sigma phosphatase inhibitor set), incubated on ice for 30 min, and centrifuged at 20,000 × g for 30 min. The supernatants were collected for western blotting.

Testis tissues were ground and re-suspended in lysis buffer, homogenized for 30 seconds with a Paddle Blender (Prima, PB100), incubated on ice for 30 min, and centrifuged at 20,000 × g for 30 min. The supernatants were collected for western blotting.

### Immunoprecipitation

The cells were cultured on 15-cm dishes and grown to confluence. Cells at 70% confluence and subjected to indicated treatment for the appropriate time according to different experiments. Then cells were washed once with PBS and harvested by scraping and centrifugation at 800 × g for 5 min. The harvested cells were washed with PBS and lysed for 30 min on ice in the lysis buffer (100 mM Tris-HCl, pH 7.4, 100 mM NaCl, 10% glycerol, 1% Triton X-100, 2 mM EDTA, Roche complete protease inhibitor set, and Sigma phosphatase inhibitor set). Cell lysates were then spun down at 12,000 × g for 20 min. The soluble fraction was collected, and the protein concentration was determined by Bradford assay. Cell extracted was mixed with anti-Flag affinity gel (Sigma-Aldrich, A2220) in a ratio of 1 mg of extract per 30 μl of agarose. After overnight rocking at 4 °C, the beads were pelleted at 2,500 × g for 3 min and washed with lysis buffer 3 times. The beads were then eluted with 0.5 mg/mL of the corresponding antigenic peptide for 6 hours or directly boiled in 1× SDS loading buffer (125 mM Tris, pH 6.8, 2% 2-mercaptoethanol, 3% SDS, 10% glycerol and 0.01% bromophenol blue).

### Immunohistochemistry and immunofluorescence

Paraffin-embedded specimens were sectioned to a 5 μm thickness and were then deparaffinized, rehydrated, and stained with haematoxylin and eosin (H&E) using standard protocols. For the preparation of the immunohistochemistry samples, sections were dewaxed, incubated in boiling citrate buffer solution for 15 min in plastic dishes, and subsequently allowed to cool down to room temperature over 3 hours. Endogenous peroxidase activity was blocked by immersing the slides in Hydrogen peroxide buffer (10%, Sinopharm Chemical Reagent) for 15 min at room temperature and were then washed with PBS. Blocking buffer (1% bovine serum albumin in PBS) was added, and the slides were incubated for 2 hours at room temperature. Primary antibody against human IgG, p-mouse-MLKL, Cleaved-caspase3, p-RIPK1, G3BP1, p-GCN2, p-PKR, p-PERK, ZBP1 or p-Human-MLKL(p-MLKL) was incubated overnight at 4°C in PBS. After 3 washes with PBS, slides were incubated with secondary antibody (polymer-horseradish-peroxidase-labeled anti-rabbit, Sigma) in PBS. After a further 3 washes, slides were analyzed using a diaminobutyric acid substrate kit (Thermo Fisher). Slides were counterstained with haematoxylin and mounted in neutral balsam medium (Sinopharm Chemical).

Immunohistochemistry analysis for SOX9/PIWIL4/DDX4/G3BP1/p-GCN2/p-PKR or p-MLKL/RIPK3 was performed using an antibody against SOX9/ PIWIL4/DDX4/G3BP1/p-GCN2/p-PKR and p-MLKL/RIPK3. Primary antibody against SOX9/PIWIL4/DDX4/G3BP1/p-GCN2/p-PKR was incubated overnight at 4°C in PBS. After 3 washes with PBS, slides were incubated with DyLight-488/555 conjugated donkey anti-rabbi/mouse secondary antibodies (Life) in PBS for 8 h at 4°C. After a further 3 washes, slides were incubated with p-MLKL/RIPK3 antibody overnight at 4°C in PBS. After a further 3 washes, slides were incubated with DyLight-488/555 conjugated donkey anti-mouse/rabbit secondary antibodies (Life) for 2 hours at room temperature in PBS. After a further 3 washes in PBS, the cell nuclei were then counterstained with DAPI (Invitrogen) in PBS. Fluorescence microscopy was performed using a Nikon A1-R confocal microscope.

Immunofluorescence analysis for G3BP1 or RIPK3 was performed using an antibody against G3BP1 and RIPK3. Primary antibody against G3BP1 was incubated overnight at 4°C in PBS. After 3 washes with PBS, slides were incubated with DyLight-555 conjugated donkey anti-rabbi/mouse secondary antibodies (Life) in PBS for 8 h at 4°C.

After a further 3 washes, slides were incubated with RIPK3 antibody overnight at 4°C in PBS. After a further 3 washes, slides were incubated with DyLight-488 conjugated donkey anti-mouse/rabbit secondary antibodies (Life) for 2 hours at room temperature in PBS. After a further 3 washes in PBS, the cell nuclei were then counterstained with DAPI (Invitrogen) in PBS. Fluorescence microscopy was performed using a Nikon A1-R confocal microscope.

### Phoshorylated MLKL peptides competition assay

Paraffin-embedded specimens were sectioned to a 5 μm thickness. Then sections were dewaxed, incubated in boiling citrate buffer solution for 15 min in plastic dishes, and subsequently allowed to cool down to room temperature over 3 hours. Endogenous peroxidase activity was blocked by immersing the slides in Hydrogen peroxide buffer (10%, Sinopharm Chemical Reagent) for 15 min at room temperature and were then washed with PBS. Blocking buffer (1% bovine serum albumin in PBS) was added, and the slides were incubated for 2 hours at room temperature. Primary antibody against p-Human-MLKL(p-MLKL) was incubated alone or co-incubates with Phoshorylated MLKL peptides overnight at 4°C in PBS. After 3 washes with PBS, slides were incubated with secondary antibody (polymer-horseradish-peroxidase-labeled anti-rabbit, Sigma) in PBS. After a further 3 washes, slides were analyzed using a diaminobutyric acid substrate kit (Thermo Fisher). Slides were counterstained with haematoxylin and mounted in neutral balsam medium (Sinopharm Chemical).

### Alkaline phosphatase dephosphorylation assay

The 5 μm paraffin sections were deparaffinized using a robotic autostainer (Leica Microsystems) and pretreated with a high pH (pH 9) buffer. After a 10-minute incubation with Peroxidase-Blocking Reagent, add 100 μL of 10X NEBuffer for Protein Metallo Phosphatases (PMP) (Cat: B0761S), 100 μL of 10 mM MnCl2 (Cat: B1761S), and 1 μL of Lambda Protein Phosphatase (Cat: P0753S, 400,000 U/mL) to make a total reaction volume of 500 μL. Incubate the samples at 37°C for 60 minutes.

Next, add Calf Intestinal Alkaline Phosphatase (CIAP) (Cat: M2825, 1,000 U/mL). Dilute CIAP in CIAP 1X Reaction Buffer to a final concentration of 500 U/mL for immediate use. Incubate the tissue sections at 37°C for 24 hours.

Following this, the samples were blocked with SuperBlock™ (TBS) Blocking Buffer for 60 minutes. The tissues were then incubated overnight in a humid chamber at room temperature (RT) with the primary p-MLKL antibody overnight at 4°C in PBS. After 3 washes with PBS, slides were incubated with secondary antibody (polymer-horseradish-peroxidase-labeled anti-rabbit, Sigma) in PBS. After a further 3 washes, slides were analyzed using a diaminobutyric acid substrate kit (Thermo Fisher). Slides were counterstained with haematoxylin and mounted in neutral balsam medium (Sinopharm Chemical).

### Quantitative RT-PCR

Cell/Tissue total RNA was extracted with the FastPure® Cell/Tissue Total RNA Isolation Kit (RC112, Vazyme Biotech, Nanjing, China) and cDNA was prepared with HiScript® III RT SuperMix for qPCR Kit (R323, Vazyme Biotech, Nanjing, China) according to the manufacturer’s protocol. Quantitative RT-PCR of ZBP1 was performed with Taq Pro Universal SYBR qPCR Master Mix (Q712, Vazyme Biotech, Nanjing, China) and the primers as follows:

ZBP1-forward: CAAGTCCTTTACCGCCTGAAG

ZBP1-reverse: TCGTCATTCCCAGAGCCTTG

GAPDH-forward: GTGCTGAGTATGTCGTGGAGTC

GAPDH-reverse: GTGGTTCACACCCATCACAAAC

Data were normalized by GAPDH expression, and relative expression changes were analyzed according to the 2^-ΔΔCT method.

### Statistical analysis

Statistical tests were used for every type of analysis. The data meet the assumptions of the statistical tests described for each figure. Results are expressed as the mean ±s.e.m or SD. Differences between experimental groups were assessed for significance using a two-tailed unpaired Student’s t-test using GraphPad prism10. The *\*P<0.05*, *\*\*P< 0.01*, \*\*\**P<0.001* and \*\*\*\**P<0.0001* levels were considered significant. P>0.05 was considered not significant (NS).

### Data availability

Source data are provided with this paper

---

**Extended Data Fig. 1.**
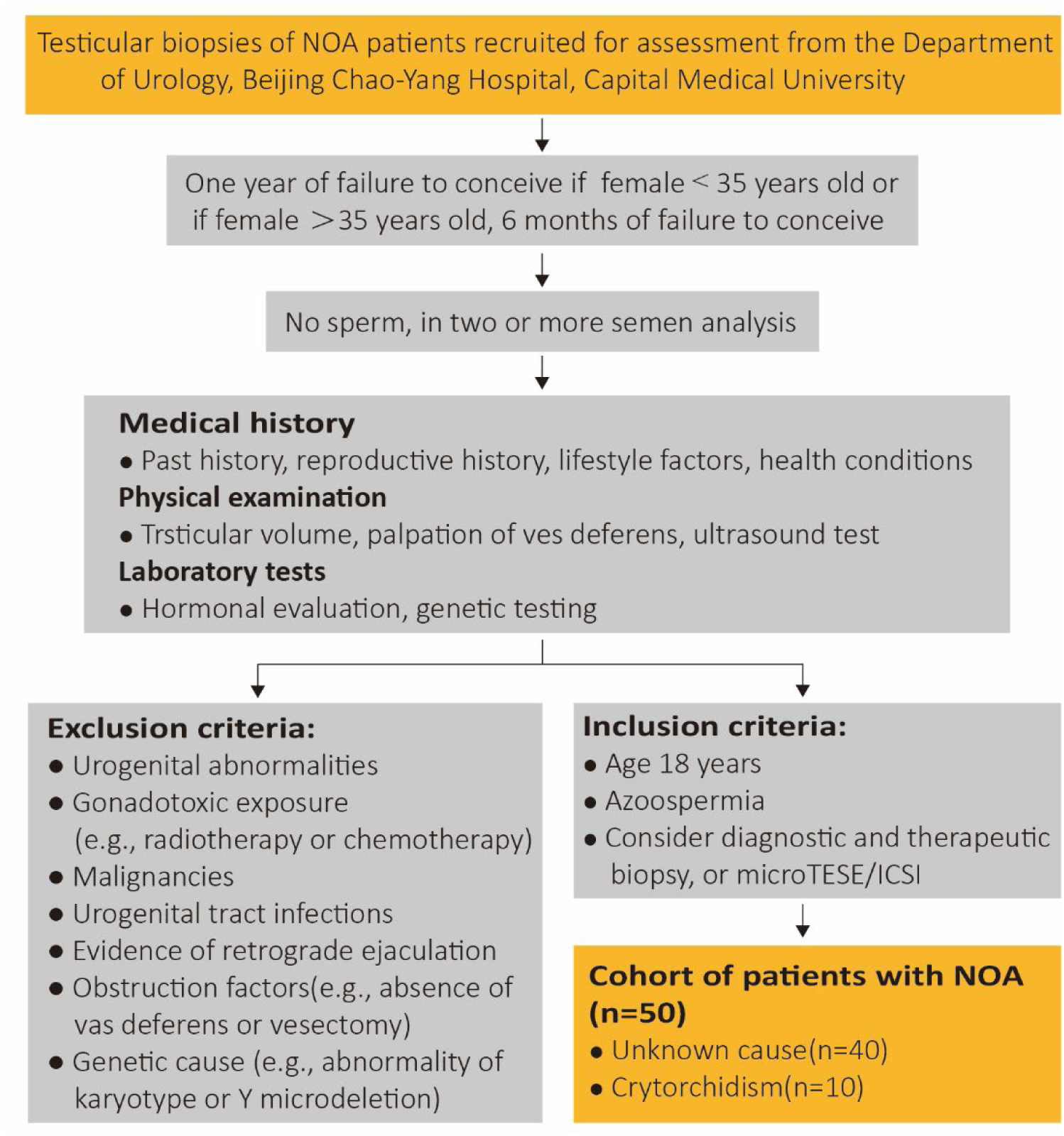
Flow chart for selecting the non-obstructive azoospermia (NOA) cohort. A total of 50 patients with NOA were recruited for the initial assessment.

**Extended Data Fig. 2.**
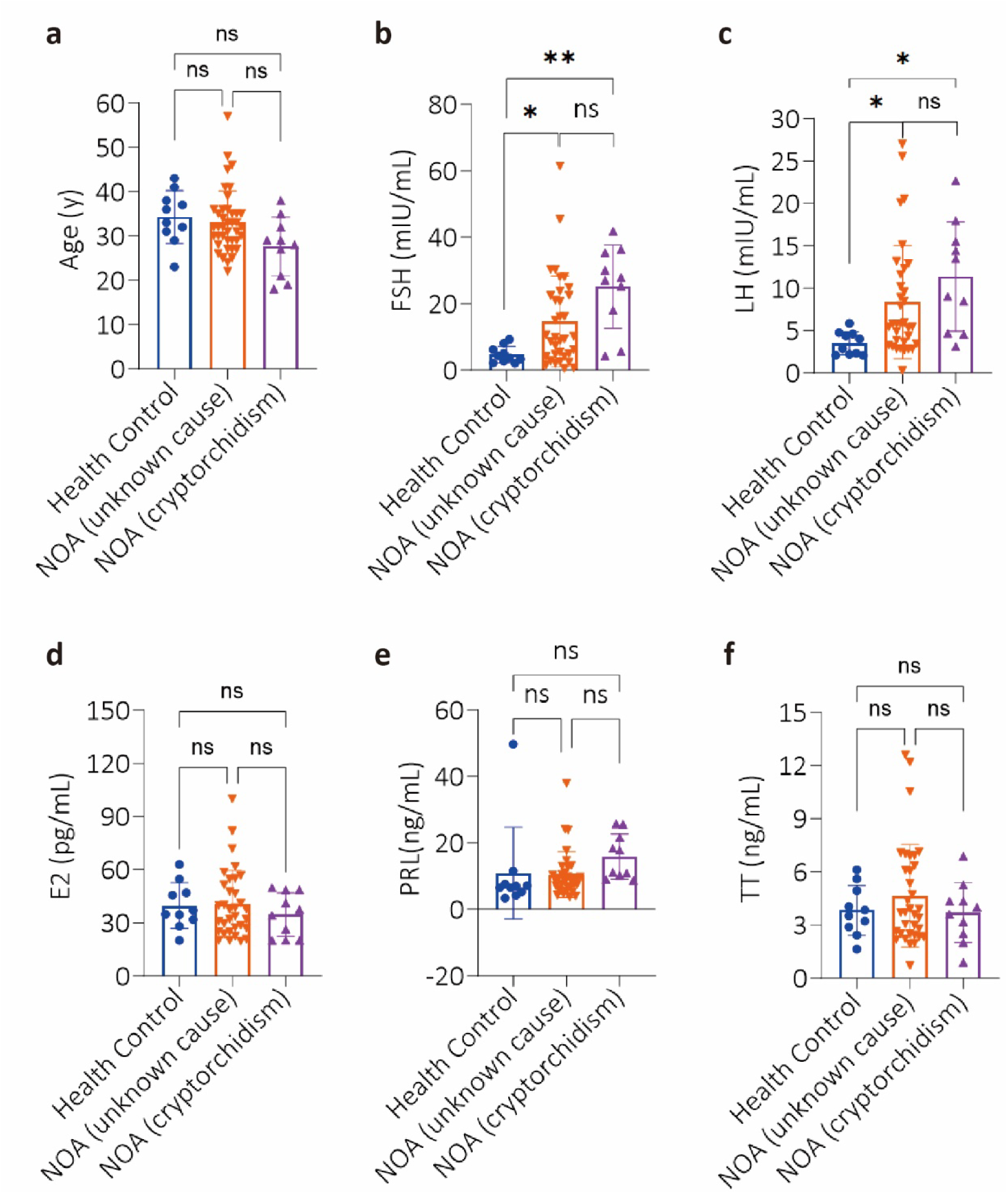
Analysis the clinical data of NOA patients. **a**, Age distribution of healthy control and NOA patients. **b**-**f**, Detecting the follicle-stimulating hormone (FSH), luteinizing hormone (LH), estradiol (E2), prolactin (PRL) and total testosterone (TT) level from healthy control(n=10) and NOA patients (unknown cause, n=40; cryptorchidism, n=10). *\*P<0.05*, *\*\*P<0.01*. *P* values were determined by two-sided unpaired Student’s *t* tests.

**Extended Data Fig. 3.**
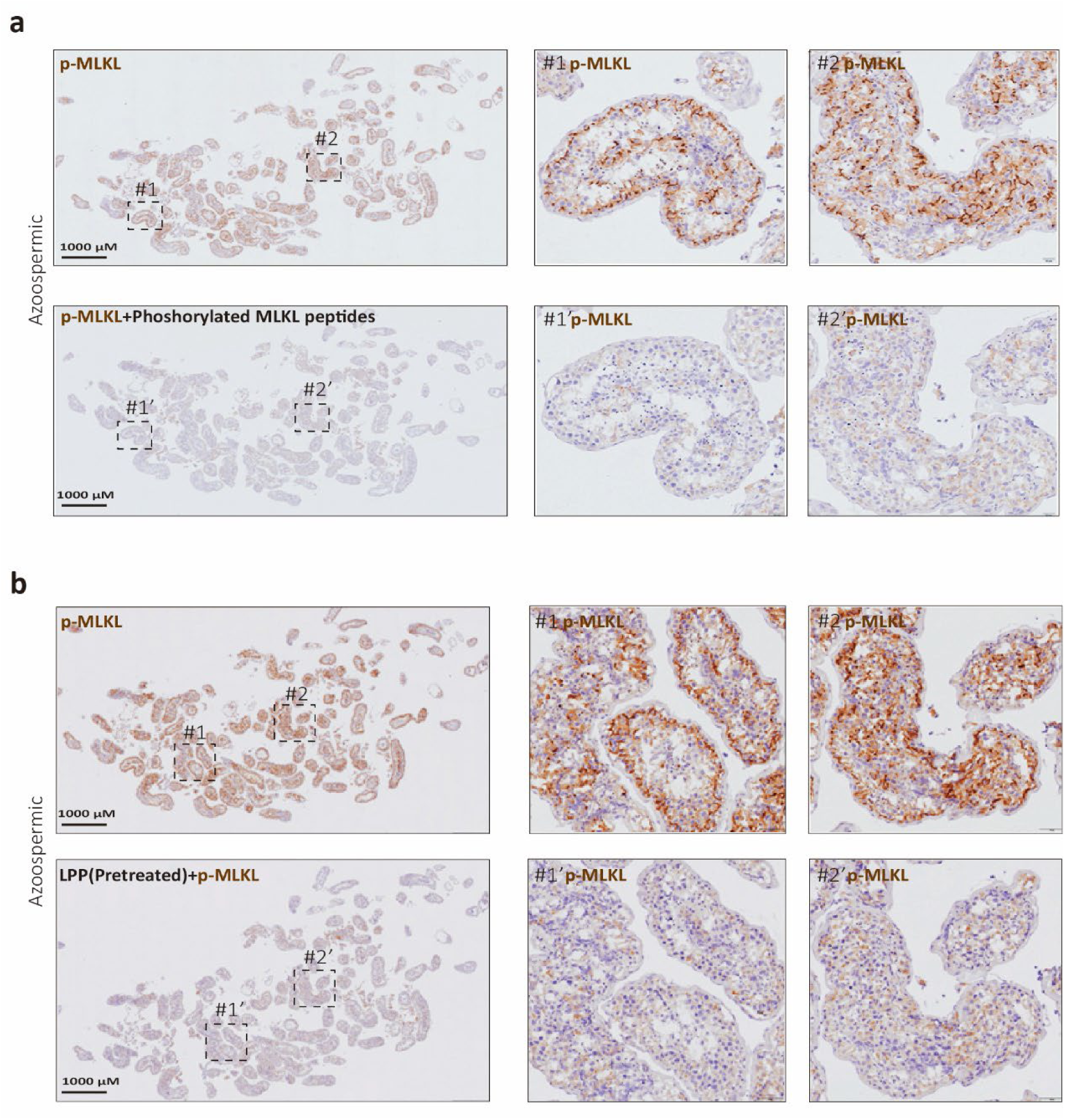
p-MLKL were detected in the seminiferous tubules of the NOA testes. **a**, Competitive IHC experiments of p-MLKL antibody. IHC analysis of NOA testes(n=5) section with p-MLKL antibody alone or co-incubated with phoshorylated MLKL (GYQVKLAGFELRKTQpTpSMSLGTTREKTDRVKS) peptides. **b**, IHC analysis of human NOA testes(n=5) selection with p-MLKL antibody with or without lambda protein phosphatase pretreatment.

**Extended Data Fig. 4.**
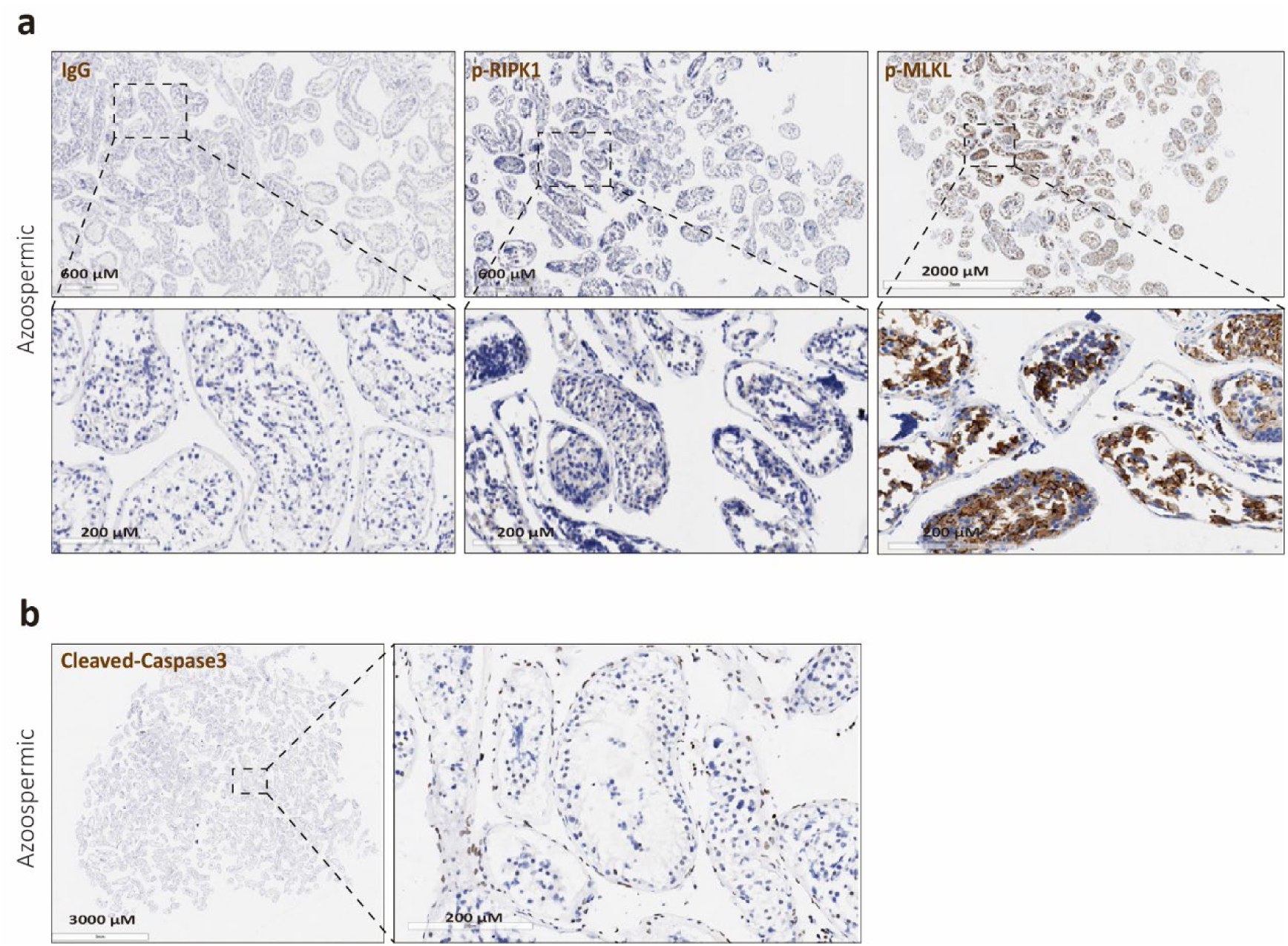
p-RIPK1 and Cleaved-caspase3 were not detected in the seminiferous tubules of the NOA testes. **a**, **b**, IHC analysis of human NOA testes (n=50) with p-RIPK1 in (a) and Cleaved-caspase3 antibodies in (b).

**Extended Data Fig. 5.**
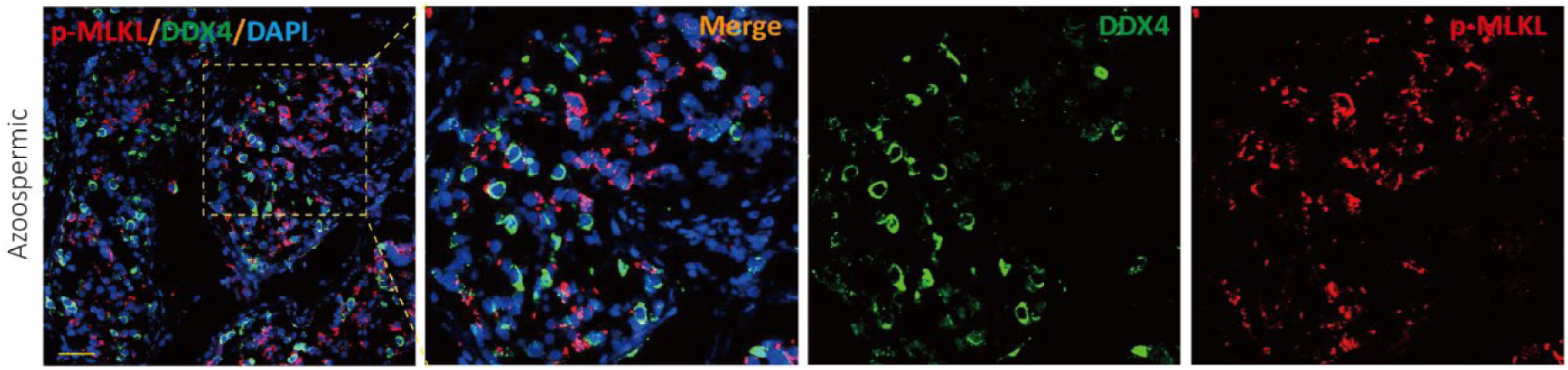
p-MLKL signal did not co-localized with DDX4 in human NOA testes selection. Immunofluorescence analysis of NOA testes (n=5) with antibodies against p-MLKL (red) and DDX4 (primary spermatocyte, secondary spermatocyte and spermatids specific protein, green). Scale bar, 50μm.

**Extended Data Fig. 6.**
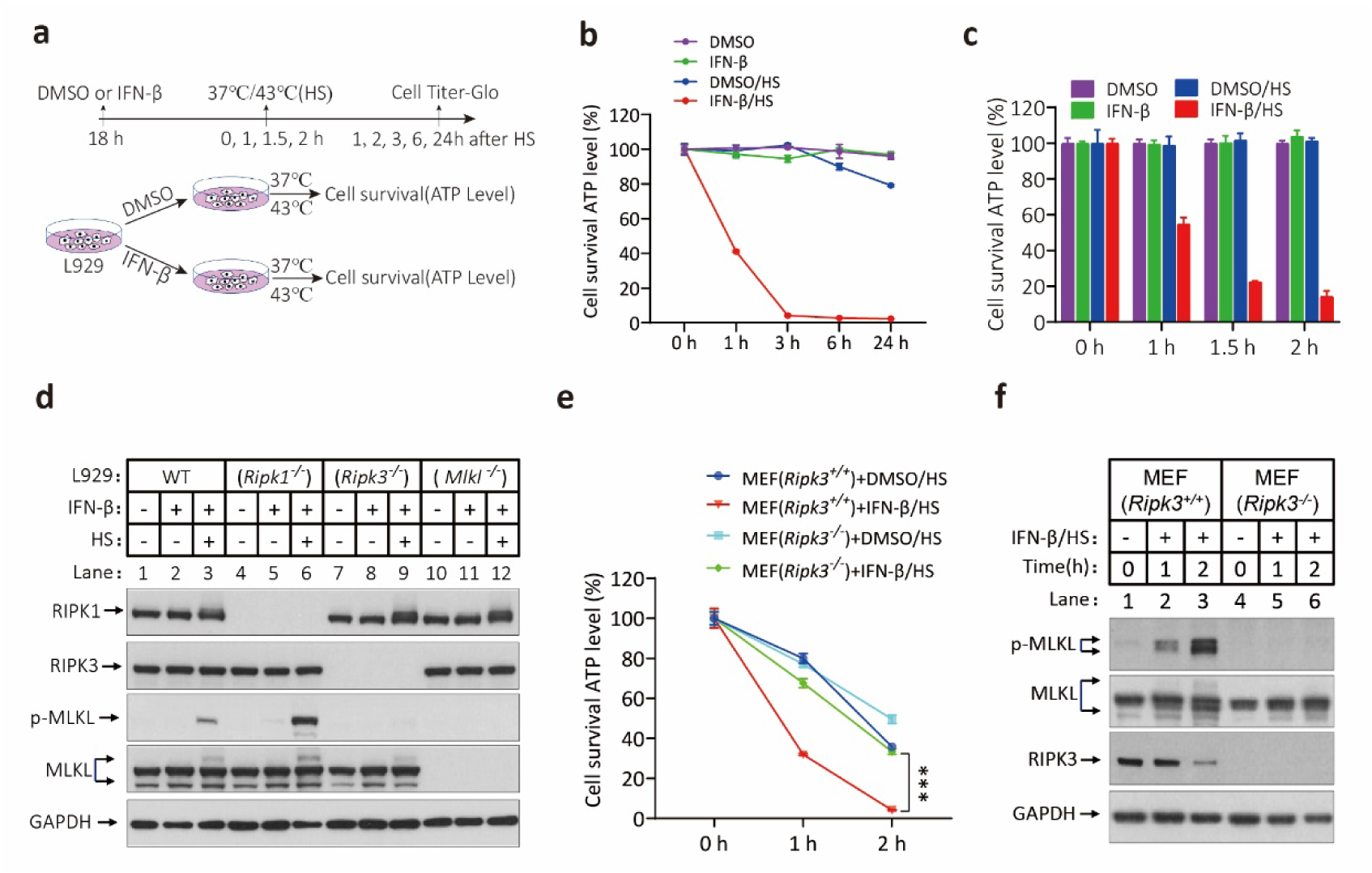
Heat shock induces RIPK3-dependent necroptosis. **a-c**, Schematic illustrating experiment design in (a). Cultured L929 cells were treated with DMSO or IFN-β for 18 hours, then the cells were transferred to 1.5 ml EP and put into 37℃ or 43℃ water for 2 hours. 0 hours,1 hours, 3 hours, 6 hours and 24 hours after heat shock cell viability as measured by Cell Titer-Glo in (b). After treatment with DMSO or IFN-β for 18 hours, L929 cells were transferred to 1.5 ml EP and put into 37℃ or 43℃ water for 0 hours, 1 hours, 1.5 hours and 2 hours. 2 hours after heat shock cell viability as measured by Cell Titer-Glo in (c). **d**, Cultured L929 cells with wild type (WT), *Ripk1*, *Ripk3* or *Mlkl* gene knocked out were treated with IFN-β and HS for 18 hours and 1.5 hours as described in Extended Data Fig. 6a. 0.5 hours after heat shock, the levels of p-MLKL, MLKL, RIPK1 and RIPK3 were analyzed by immunoblotting, GAPDH was used as loading control. **e**, **f**, Cultured wild type MEF, or MEF with their *Ripk3* gene knocked out were treated with DMSO or IFNβ/HS as indicated. The cell viability was measured by Cell-titer Glo in (e). The levels of p-MLKL, MLKL and RIPK3 were analyzed by immunoblotting in (f), GAPDH was used as loading control. Data in (b, c and e) represent the mean ± SD. \*\*\**P*<0.001. *P* values were determined by two-sided unpaired Student’s *t* tests.

**Extended Data Fig. 7.**
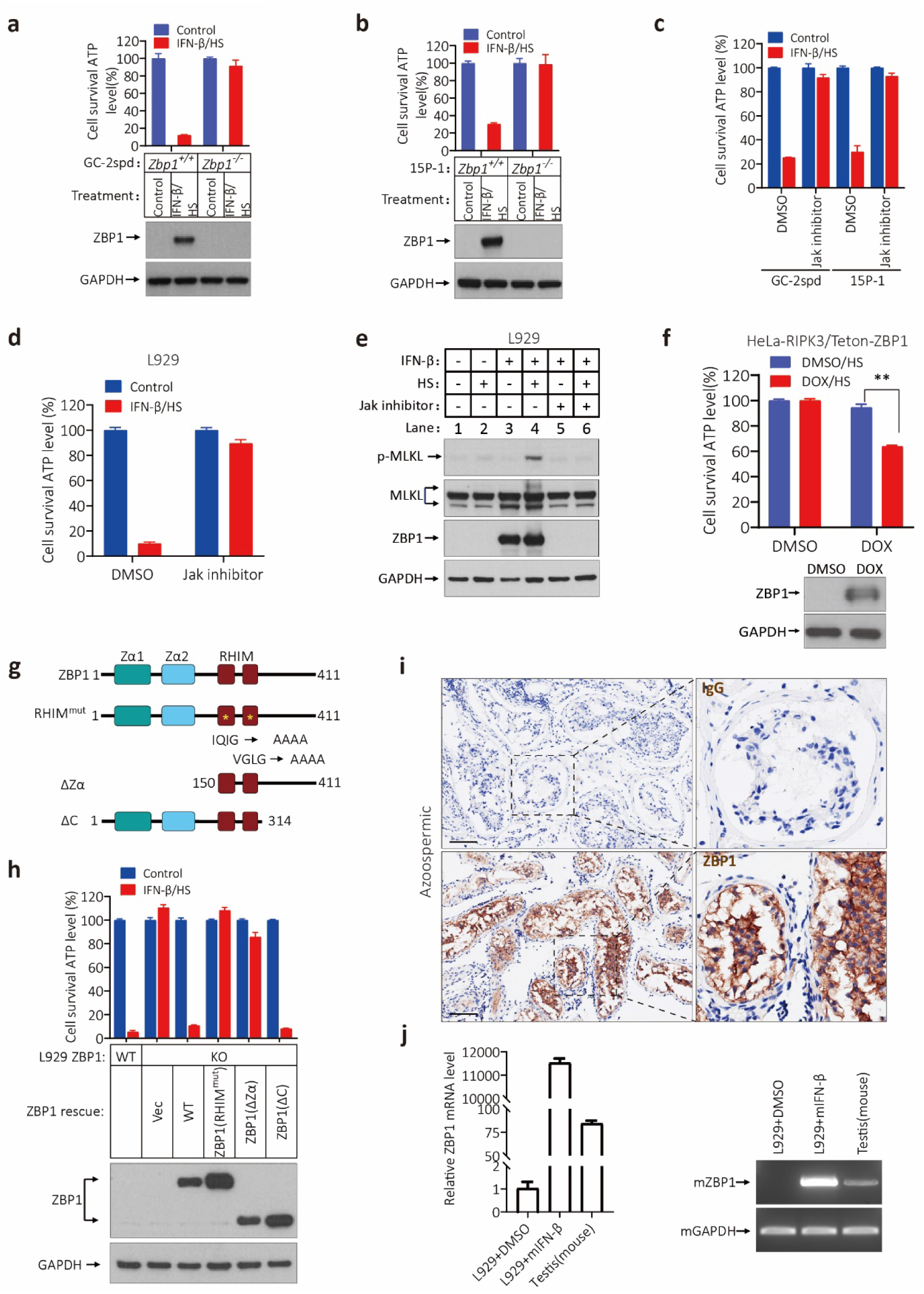
Heat shock-induced RIPK3-dependent necroptosis requires Jak/STAT-mediated ZBP1 expression. **a**, **b**, Cultured GC-2spd and 15P-1 cells with wild type or *Zbp1* gene knocked out were treated with IFN-β and HS for 18 hours and 1.5 hours as described in Fig. 2a, 6 hours after heat shock cell viability as measured by Cell Titer-Glo in (a, b). The level of ZBP1 was analyzed by immunoblotting, GAPDH was used as loading control. **c**-**e**, Cultured GC-2spd, 15P-1 and L929 cells were treated with DMSO, IFN-β, Jak inhibitor or IFN-β+Jak inhibitor for 18 hours, then the cells were transferred to 1.5 ml EP and put into 43℃ water for 1.5 hours, 6 or 2 hours after heat shock cell viability as measured by Cell Titer-Glo in (c, d). The levels of p-MLKL, MLKL and ZBP1 were analyzed by immunoblotting in (e), GAPDH was used as loading control. **f**, Cultured HeLa-RIPK3/Teton-ZBP1 cells were treated with DMSO/HS or DOX/HS as indicated. The cell viability was measured by Cell-titer Glo. The levels of ZBP1 were analyzed by immunoblotting, GAPDH was used as loading control. **g**, **h**, Schematic representation of full-length ZBP1 or indicated mutants in (g). Cultured L929 wild type, or L929 with their *Zbp1* gene knocked out cells were infected with lentivirus expressing vector control (Vec), wild type ZBP1 (WT) or its truncation mutants followed by treatment of stimuli as indicated. The cell viability was measured by Cell-titer Glo in (h). The levels of ZBP1 and its truncation mutants were analyzed by immunoblotting in (h). **i**, IHC analysis of human NOA testes (n=3) with IgG and ZBP1 antibodies. Scale bar, 100 μm. **j**, Quantitative RT-PCR analysis (mean ± SD, n=3) of mouse ZBP1 in wild type adult testis and L929 cells after INF-β treatment for 18 hours. The PCR products were analyzed by agarose gel electrophoresis. Data in (a, b, c, d, f and h) represent the mean ± SD \*\**P*<0.01. *P* values were determined by two-sided unpaired Student’s *t* tests.

**Extended Data Fig. 8.**
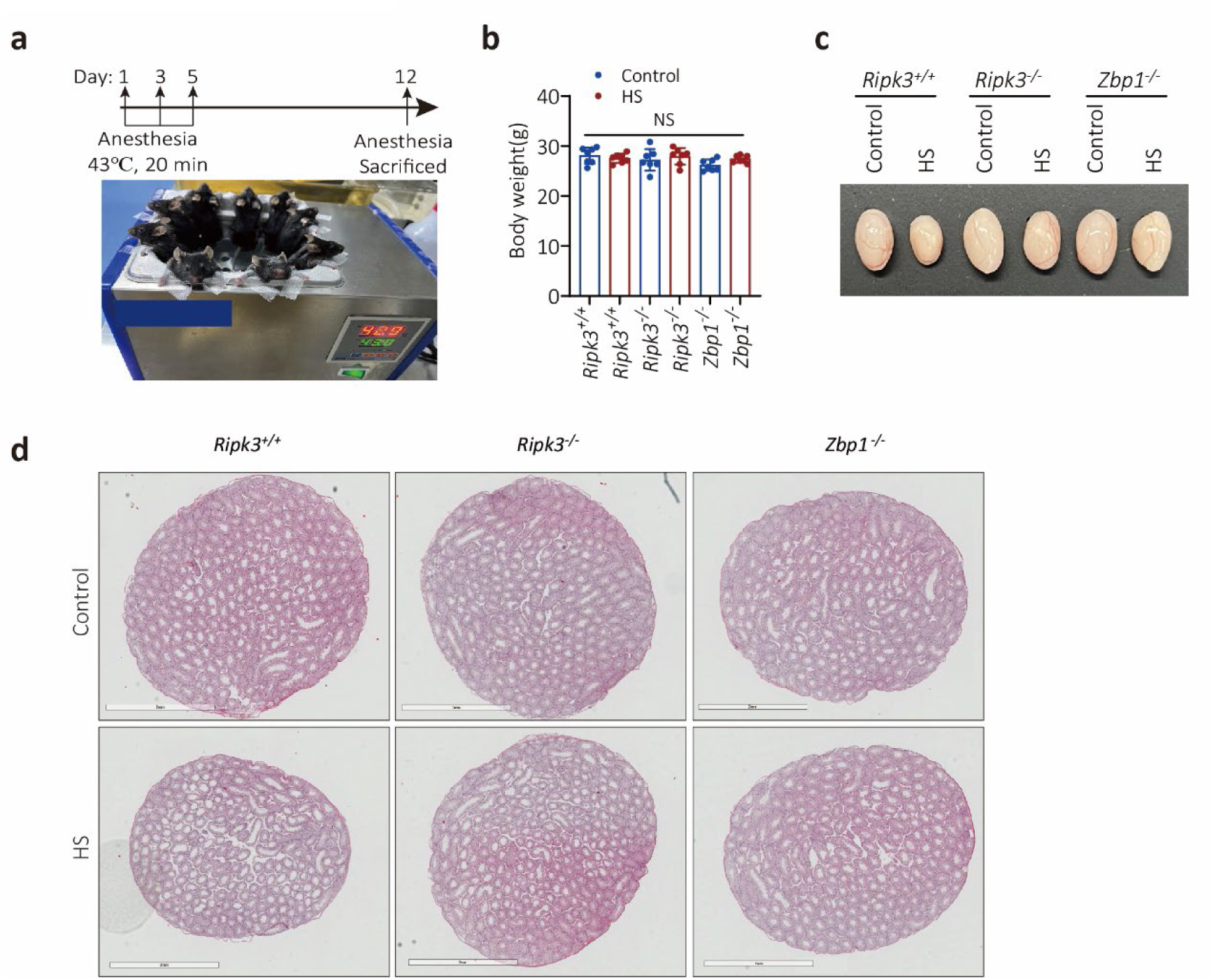
Rescuing the heat shock-induced testes damage with *Ripk3* or *Zbp1* Knockout. **a**, Schematic illustrating experiment design in (a), the lower body of mice were put in 43℃ water for 20 min every other day for 3 times, 7 days after heat shock, mice were sacrificed and testes were weighted. The pictures of mice under heat shock shown below. **b**, The weights of whole body of 12-week-old *Ripk3^+/+^*, *Ripk3^-/-^*littermate and *Zbp1^-/-^* male mice (n=7 for each genotype) after heat shock treatment for 7 days. Data represent the mean ± s.e.m. *P* values were determined by two-sided unpaired Student’s *t* tests. Not significant (NS). **c**, Macroscopic features of testes of 12-week-old *Ripk3^+/+^*, *Ripk3^-/-^* littermate and *Zbp1^-/-^* male mice after heat shock treatment for 7 days. **d**, H&E staining of testis sections from 12-week-old *Ripk3^+/+^*, *Ripk3^-/-^* littermate and *Zbp1^-/-^* male mice after heat shock treatment for 7 days. Scale bar, 2 mm.

**Extended Data Fig. 9.**
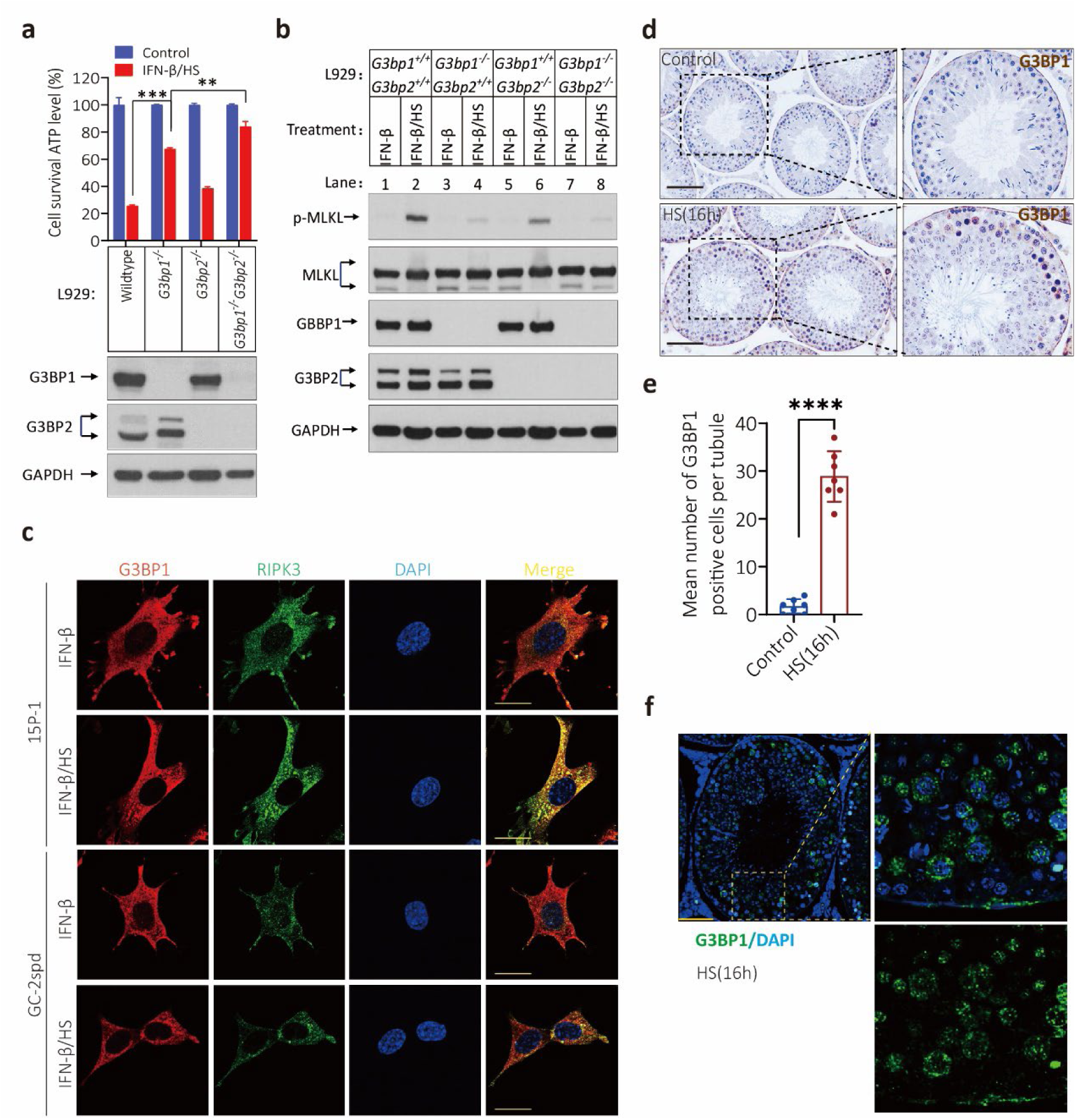
Stress granule is required for heat shock-induced necroptosis. **a**, **b**, Cultured L929, L929(*G3bp1^-/-^*), L929(*G3bp2^-/-^*) and L929(*G3bp1^-/-^G3bp2^-/-^*) cells were treated with IFN-β and HS for 18 hours and 1.5 hours as described in Extended Data Fig. 6a, the intracellular ATP levels were measured by Cell Titer-Glo in (a). The levels of p-MLKL, MLKL, ZBP1, G3BP1, and G3BP2 were analyzed by immunoblotting in (b), GAPDH was used as loading control. Data in (a) are mean ± SD of triplicate wells. \*\**P*<0.01. \*\*\**P*<0.001. *P* values were determined by two-sided unpaired Student’s *t* tests. **c**, Cultured GC-2spd and 15P-1 cells were first treated with the indicated stimuli for 18h(IFN-β), then heat shock for 30 min. The RIPK3 and G3BP1 were detected by immunofluorescence. Scale bares, 10 μm. **d**, **e**, IHC staining of testes from 12-week-old wildtype male littermate mice (n=7 for each genotype) with or without heat shock for 16 hours with an anti-G3BP1 antibody in (d). G3BP1 positive cells were counted in five fields per testis and quantified in (e). Scale bar, 100 μm. Data represent the mean ± s.e.m. \*\*\*\**P*<0.001. *P* values were determined by two-sided unpaired Student’s *t* tests. **f**, Immunofluorescence analysis of testes after heat shock for 16 hours(n=7) with antibodies against G3BP1(green). Scale bar, 50 μm.

**Extended Data Fig. 10.**
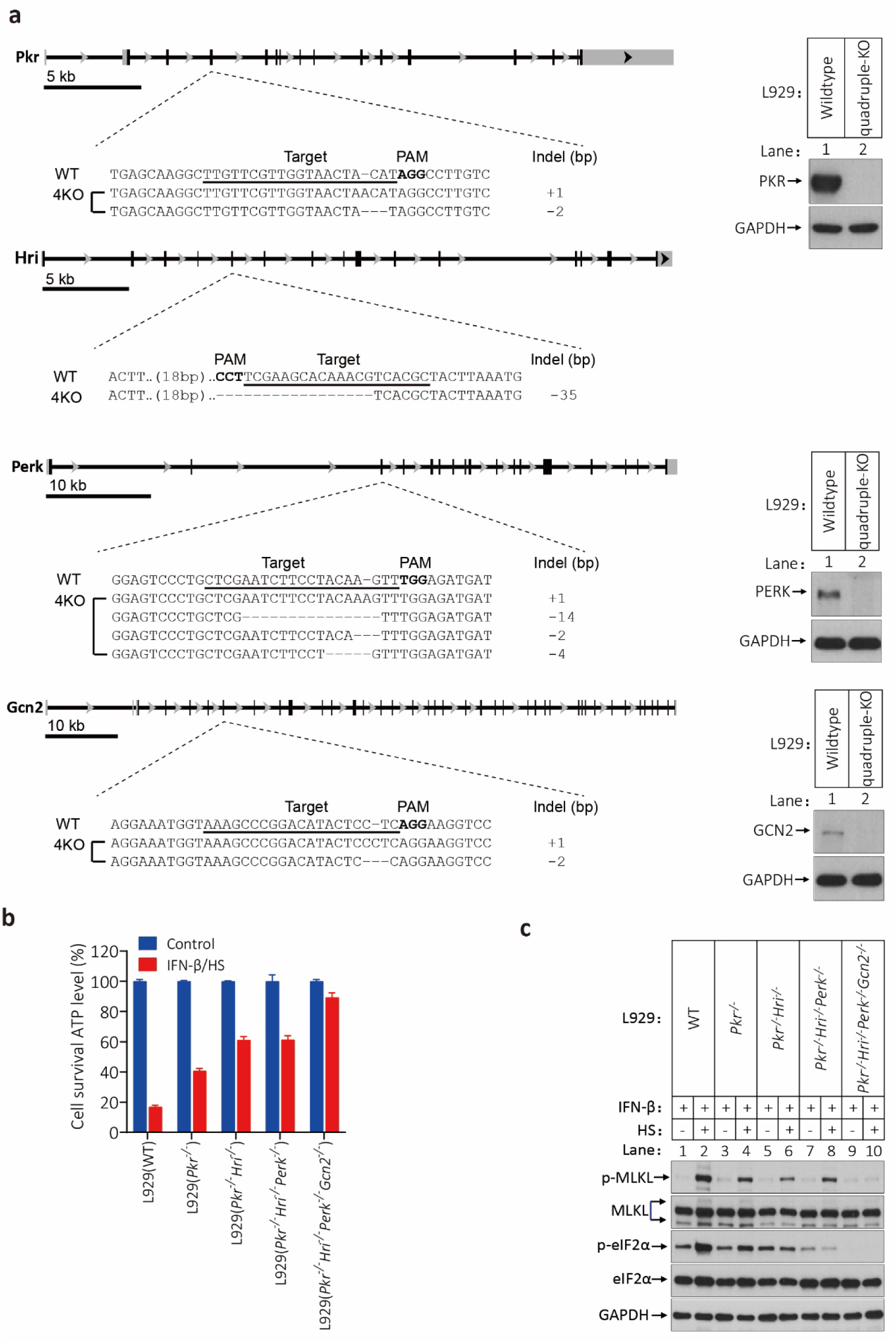
*Eif2α* kinases are required for heat shock-induced necroptosis. **a**, Schematic of each targeting locus of the four *Eif2α* kinase with the target sequence underscored, the guide RNA sequences targeting the exon of each gene was shown with the PAM sequences highlighted in bold type. Except HRI, the other three kinases protein levels were analyzed by immunoblotting. **b**, **c**, Cultured L929 cells with wild type, signal-knockout (*Pkr* knockout) double-knockout (*Pkr* and *Hri* knockout), triple-knockout (*Pkr*, *Hri* and *Perk* knockout) and quadruple-knockout (*Pkr*, *Hri*, *Perk* and *Gcn2* knockout) were treated with IFN-β and HS for 18 hours and 1.5 hours as described in Extended Data Fig. 6a, the intracellular ATP levels were measured by Cell Titer-Glo in (b). The levels of p-MLKL, MLKL, eIF2α and p-eIF2α were analyzed by immunoblotting in (c), GAPDH was used as loading control. Data in (b) are mean ± SD of triplicate wells.

**Extended Data Fig. 11.**
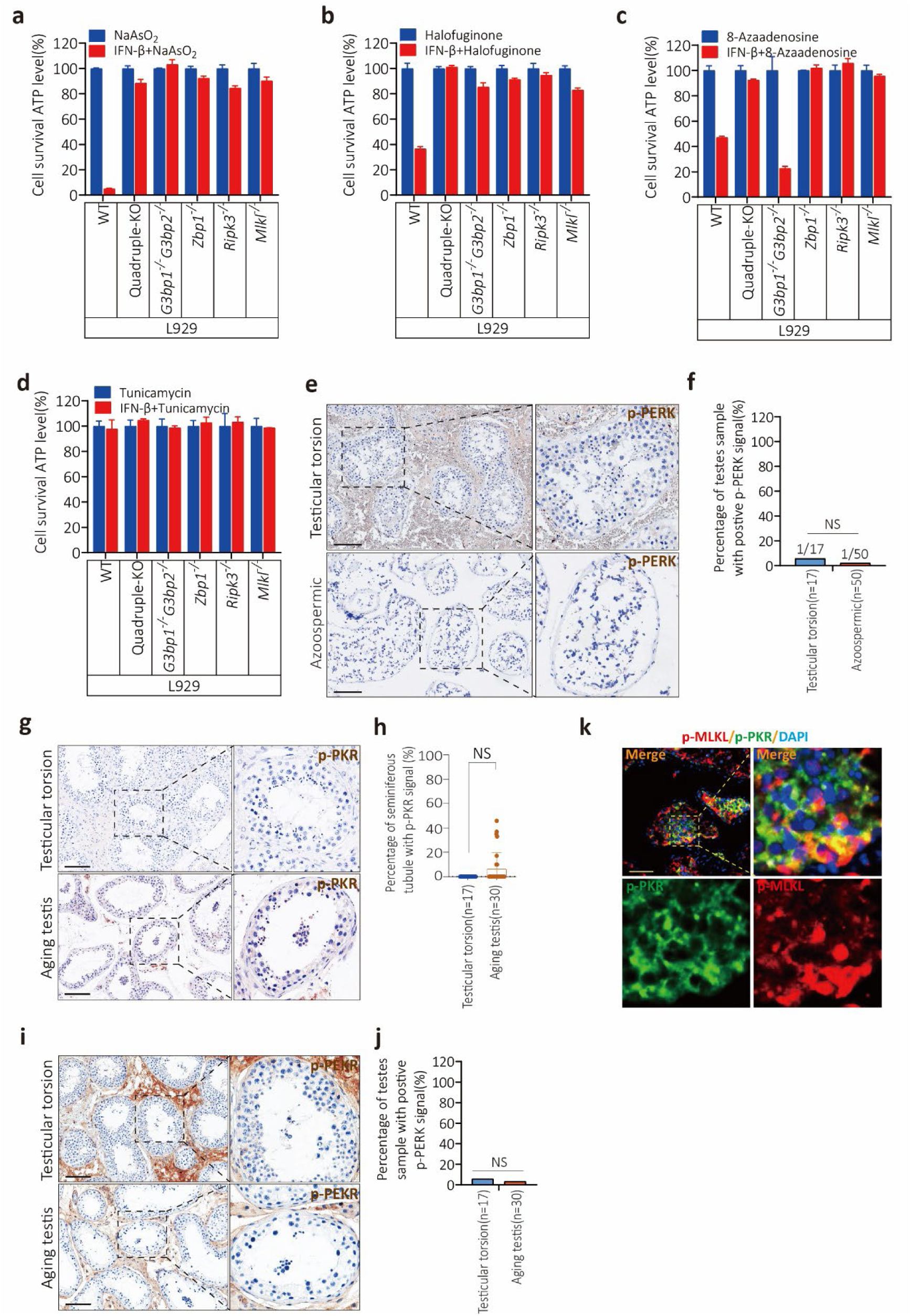
The triggers of stress induce ZBP1 and RIPK3-dependent necroptosis in L929 cells. a-d. , Cultured L929, L929(*Pkr^-/-^Hri^-/-^Perk^-/-^Gcn2^-/-^*), L929(*G3bp1^-/-^G3bp2^-/-^*), L929(*Zbp1^-/-^*), L929(*Ripk3^-/-^*) and L929(*Mlkl^-/-^*) cells were treated with NaAsO_2_ (Sodium arsenite, 30 μM) and NaAsO_2_(30 μM)+IFN-β(10 ng/ml) for 18 hours; Halofuginone(500 nM) and Halofuginone(500 nM)+IFN-β(10 ng/ml) for 42 hours; 8-Azaadenosine (20 μM) and 8-Azaadenosine(20 μM)+IFN-β(10 ng/ml) for 46 hours; Tunicamycin (2.5 μM) and Tunicamycin(2.5 μM)+IFN-β(10 ng/ml) for 30 hours. The intracellular ATP levels were measured by Cell Titer-Glo in (a-d). Data are mean ± SD of triplicate wells. **e**, **f**, Immunohistochemistry analysis of human testicular torsion and NOA testes with p-PERK antibody in (e). Scale bar, 100 μm. The number of seminiferous tubules with positive p-PERK signal were counted based on IHC staining and quantification in (f). *P* values were determined by two-sided unpaired Student’s *t* tests. Not significant (NS). **g**, **h**, IHC analysis of human testicular torsion and aging testis sections with p-PKR antibody in (g). The number of seminiferous tubules with positive p-PKR signal were counted based on IHC staining and quantification in (h). Scale bar, 100 μm. Data represent the mean ± s.e.m. *P* values were determined by two-sided unpaired Student’s t tests. Not significant (NS). **i**, **j**, IHC analysis of human testicular torsion and aging testis sections with p-PEKR antibody in (i). The number of seminiferous tubules with positive p-PEKR signal were counted based on IHC staining and quantification in (j). Scale bar, 100 μm. Data represent the mean ± s.e.m. *P* values were determined by two-sided unpaired Student’s t tests. Not significant (NS). **k**, Immunofluorescence analysis of aging testis sections(n=5) with antibodies against p-MLKL (red) and p-PKR (green). Scale bar, 50 μm.

**Extended Data Fig. 12.**
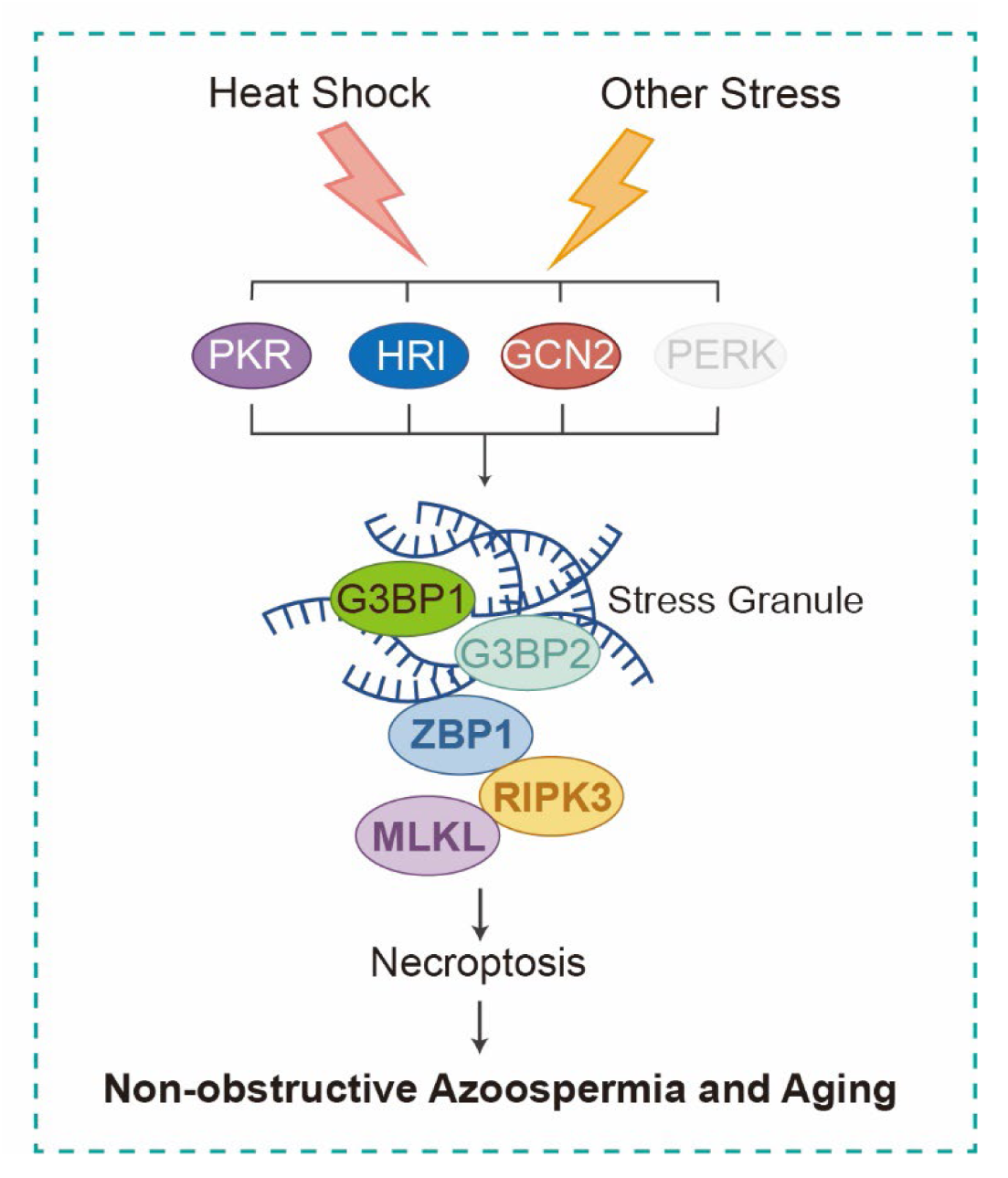
Stress granule induces ZBP1-RIPK3-dependent necroptosis leading to non-obstructive azoospermia and testis aging. Heat shock or other stress activates eIF2α kinases (PKR, HRI or GCN2). The activated stress kinase promotes stress granule formation and recruits ZBP1 to form ZBP1-RIPK3 necrosomes, then RIPK3 will be activated. Activated RIPK3 phosphorylates MLKL and executes necroptosis in testes cells leading to non-obstructive azoospermia and testis aging.

**Supplementary Table 1.** The clinical characteristics of health control(n=10), testicular torsion patients (n=17), NOA patients (unknown cause, n=40; cryptorchidism, n=10) and prostate cancer patients (n=30).

## References

1 Agarwal, A., Mulgund, A., Hamada, A. & Chyatte, M. R. A unique view on male infertility around the globe. Reprod Biol Endocrinol 13, 37, doi:10.1186/s12958-015-0032-1 (2015).

2 Willott, G. M. Frequency of azoospermia. Forensic Sci Int 20, 9–10, doi:10.1016/0379-0738(82)90099-8 (1982).

3 McLachlan, R. I., Rajpert-De Meyts, E., Hoei-Hansen, C. E., de Kretser, D. M. & Skakkebaek, N. E. Histological evaluation of the human testis--approaches to optimizing the clinical value of the assessment: mini review. Hum Reprod 22, 2–16, doi:10.1093/humrep/del279 (2007).

4 Agarwal, A. et al. Male infertility. Lancet 397, 319–333, doi:10.1016/S0140-6736(20)32667-2 (2021).

5 Jimbo, M. et al. Fertility in the aging male: a systematic review. Fertil Steril 118, 1022–1034, doi:10.1016/j.fertnstert.2022.10.035 (2022).

6 Pohl, E., Gromoll, J., Wistuba, J. & Laurentino, S. Healthy ageing and spermatogenesis. Reproduction 161, R89–R101, doi:10.1530/REP-20-0633 (2021).

7 Nie, X. et al. Single-cell analysis of human testis aging and correlation with elevated body mass index. Dev Cell 57, 1160–1176 e1165, doi:10.1016/j.devcel.2022.04.004 (2022).

8 He, S. & Wang, X. RIP kinases as modulators of inflammation and immunity. Nat Immunol 19, 912–922, doi:10.1038/s41590-018-0188-x (2018).

9 Pasparakis, M. & Vandenabeele, P. Necroptosis and its role in inflammation. Nature 517, 311–320, doi:10.1038/nature14191 (2015).

10 Ai, Y., Meng, Y., Yan, B., Zhou, Q. & Wang, X. The biochemical pathways of apoptotic, necroptotic, pyroptotic, and ferroptotic cell death. Mol Cell 84, 170–179, doi:10.1016/j.molcel.2023.11.040 (2024).

11 Li, D. et al. RIPK1-RIPK3-MLKL-dependent necrosis promotes the aging of mouse male reproductive system. Elife 6, doi:10.7554/eLife.27692 (2017).

12 Li, D. et al. Casein kinase 1G2 suppresses necroptosis-promoted testis aging by inhibiting receptor-interacting kinase 3. Elife 9, doi:10.7554/eLife.61564 (2020).

13 Brannigan, R. E. et al. Updates to Male Infertility: AUA/ASRM Guideline (2024). J Urol 212, 789–799, doi:10.1097/JU.0000000000004180 (2024).

14 Xie, Y. et al. Phosphorylated mixed lineage kinase domain-like protein in human seminal plasma: A potential novel biomarker of spermatogenic function. Andrologia 51, e13310, doi:10.1111/and.13310 (2019).

15 Sun, L. et al. Mixed lineage kinase domain-like protein mediates necrosis signaling downstream of RIP3 kinase. Cell 148, 213–227, doi:10.1016/j.cell.2011.11.031 (2012).

16 De Toni, L., Finocchi, F., Jawich, K. & Ferlin, A. Global warming and testis function: A challenging crosstalk in an equally challenging environmental scenario. Front Cell Dev Biol 10, 1104326, doi:10.3389/fcell.2022.1104326 (2022).

17 Jorban, A., Lunenfeld, E. & Huleihel, M. Effect of Temperature on the Development of Stages of Spermatogenesis and the Functionality of Sertoli Cells In Vitro. Int J Mol Sci 25, doi:10.3390/ijms25042160 (2024).

18 Yuan, F. et al. Z-DNA binding protein 1 promotes heatstroke-induced cell death. Science 376, 609–615, doi:10.1126/science.abg5251 (2022).

19 Ingram, J. P. et al. ZBP1/DAI Drives RIPK3-Mediated Cell Death Induced by IFNs in the Absence of RIPK1. Journal of immunology (Baltimore, Md. : 1950) 203, 1348–1355, doi:10.4049/jimmunol.1900216 (2019).

20 Yang, D. et al. ZBP1 mediates interferon-induced necroptosis. Cell Mol Immunol 17, 356–368, doi:10.1038/s41423-019-0237-x (2020).

21 Kuriakose, T. & Kanneganti, T. D. ZBP1: Innate Sensor Regulating Cell Death and Inflammation. Trends Immunol 39, 123–134, doi:10.1016/j.it.2017.11.002 (2018).

22 Hasani, A. et al. Non-apoptotic cell death such as pyroptosis, autophagy, necroptosis and ferroptosis acts as partners to induce testicular cell death after scrotal hyperthermia in mice. Andrologia 54, e14320, doi:10.1111/and.14320 (2022).

23 Ziaeipour, S. et al. Hyperthermia versus busulfan: Finding the effective method in animal model of azoospermia induction. Andrologia 51, e13438, doi:10.1111/and.13438 (2019).

24 Zhang, P. et al. Melatonin protects the mouse testis against heat-induced damage. Mol Hum Reprod 26, 65–79, doi:10.1093/molehr/gaaa002 (2020).

25 Protter, D. S. W. & Parker, R. Principles and Properties of Stress Granules. Trends Cell Biol 26, 668–679, doi:10.1016/j.tcb.2016.05.004 (2016).

26 Glauninger, H., Wong Hickernell, C. J., Bard, J. A. M. & Drummond, D. A. Stressful steps: Progress and challenges in understanding stress-induced mRNA condensation and accumulation in stress granules. Mol Cell 82, 2544–2556, doi:10.1016/j.molcel.2022.05.014 (2022).

27 Kedersha, N. L., Gupta, M., Li, W., Miller, I. & Anderson, P. RNA-binding proteins TIA-1 and TIAR link the phosphorylation of eIF-2 alpha to the assembly of mammalian stress granules. J Cell Biol 147, 1431–1442, doi:10.1083/jcb.147.7.1431 (1999).

28 Solomon, S. et al. Distinct structural features of caprin-1 mediate its interaction with G3BP-1 and its induction of phosphorylation of eukaryotic translation initiation factor 2alpha, entry to cytoplasmic stress granules, and selective interaction with a subset of mRNAs. Mol Cell Biol 27, 2324–2342, doi:10.1128/MCB.02300-06 (2007).

29 Arimoto, K., Fukuda, H., Imajoh-Ohmi, S., Saito, H. & Takekawa, M. Formation of stress granules inhibits apoptosis by suppressing stress-responsive MAPK pathways. Nat Cell Biol 10, 1324–1332, doi:10.1038/ncb1791 (2008).

30 Szczerba, M. et al. Canonical cellular stress granules are required for arsenite-induced necroptosis mediated by Z-DNA-binding protein 1. Sci Signal 16, eabq0837, doi:10.1126/scisignal.abq0837 (2023).

31 Pakos-Zebrucka, K. et al. The integrated stress response. EMBO Rep 17, 1374–1395, doi:10.15252/embr.201642195 (2016).

32 Holcik, M. & Sonenberg, N. Translational control in stress and apoptosis. Nat Rev Mol Cell Biol 6, 318–327, doi:10.1038/nrm1618 (2005).

33 Taniuchi, S., Miyake, M., Tsugawa, K., Oyadomari, M. & Oyadomari, S. Integrated stress response of vertebrates is regulated by four eIF2alpha kinases. Sci Rep 6, 32886, doi:10.1038/srep32886 (2016).

